# Highly basic clusters in the HSV-1 nuclear egress complex drive membrane budding by inducing lipid ordering

**DOI:** 10.1101/2021.05.18.444627

**Authors:** Michael K. Thorsen, Alex Lai, Michelle W. Lee, David P. Hoogerheide, Gerard C. L. Wong, Jack H. Freed, Ekaterina E. Heldwein

## Abstract

During replication of herpesviruses, capsids escape from the nucleus into the cytoplasm by budding at the inner nuclear membrane. This unusual process is mediated by the viral nuclear egress complex (NEC) that deforms the membrane around the capsid by oligomerizing into a hexagonal, membrane-bound scaffold. Here, we found that highly basic membrane-proximal regions (MPRs) of the NEC alter lipid order by inserting into the lipid headgroups and also promote negative Gaussian curvature. We also find that the electrostatic interactions between the MPRs and the membranes are essential for membrane deformation. One of the MPRs is phosphorylated by a viral kinase during infection, and the corresponding phosphomimicking mutations block capsid nuclear egress. We show that the same phosphomimicking mutations disrupt the NEC/membrane interactions and inhibit NEC-mediated budding *in vitro*, providing a biophysical explanation for the *in-vivo* phenomenon. Our data suggest that the NEC generates negative membrane curvature by both lipid ordering and protein scaffolding and that phosphorylation acts as an “off” switch that inhibits the membrane-budding activity of the NEC to prevent capsid-less budding.

## Introduction

To overcome the barriers presented by compartmentalization in eukaryotic cells, viruses must manipulate cellular membranes. One of the more unusual mechanisms of membrane remodeling is found in herpesviruses – large double-stranded-DNA viruses that infect nearly all vertebrates and some invertebrates for life (Beurden & Engelsma, 2012) and in humans can cause symptoms ranging from painful skin lesions to blindness and life-threatening conditions in people with weak or immature immune systems (Roizman, 2013). After viral genomes are replicated and packaged, herpesviral capsids have to traverse several host membrane barriers to complete their assembly and exit the cell as infectious virions (reviewed in (Bigalke & Heldwein, 2017; Draganova *et al*, 2020; Johnson & Baines, 2011; Mettenleiter *et al*, 2009)). The critical, conserved first step in this process is nuclear egress, during which newly formed capsids translocate from the nucleus into the cytoplasm. Many viruses that replicate their genomes within the nucleus, such as HIV, influenza, and polyomaviruses, exit this double-membraned organelle through nuclear pores (Flint *et al*, 2015; Whittaker & Helenius, 1998). However, the ~50-nm opening of the nuclear pore is too small to accommodate the ~125-nm herpesviral capsids. Instead, herpesviruses use a different, non-canonical nuclear export mechanism where capsids acquire envelopes by budding at the inner nuclear membrane (INM) and pinching off into the perinuclear space. These perinuclear enveloped virions then fuse with the outer nuclear membrane (ONM), releasing the capsids into the cytoplasm (reviewed in (Bigalke & Heldwein, 2017; Draganova *et al.*, 2020; Johnson &Baines, 2011; Mettenleiter, 2016; Roller & Baines, 2017)).

Capsid budding at the INM requires the generation of negative membrane curvature by the viral nuclear egress complex (NEC), a heterodimer of two conserved viral proteins: UL31, a soluble nuclear phosphoprotein, and UL34, which contains a single C-terminal transmembrane (TM) helix that anchors the NEC in the INM (reviewed in (Draganova *et al.*, 2020)). Both UL31 and UL34 are essential for nuclear egress, and in the absence of either protein, capsids accumulate in the nucleus and the production of infectious virions is significantly impaired (Bubeck *et al*, 2004; Chang *et al*, 1997; Farina *et al*, 2005; Fuchs *et al*, 2002; Granato *et al*, 2008; Haugo *et al*, 2011; Klupp *et al*, 2000; Lötzerich *et al*, 2006; Neubauer *et al*, 2002; Roller *et al*, 2000). Using *in-vitro* model systems and cryogenic electron microscopy and tomography (cryoEM/ET), we previously discovered that the NEC from a prototypical herpes simplex virus type 1 (HSV-1) vesiculates synthetic lipid bilayers *in vitro* in the absence of any other factors or ATP (Bigalke *et al*, 2014), which was later confirmed with the NEC homolog from a closely related pseudorabies virus (PRV) (Lorenz *et al*, 2015b). Likewise, overexpression of PRV or Epstein-Barr virus (EBV) NEC in uninfected cells caused formation of capsidless vesicles in the perinuclear space (Desai *et al*, 2012; Klupp *et al*, 2007). Furthermore, cryoEM studies showed that the NEC oligomerizes into hexagonal scaffold-like coats on the inner surface of budded vesicles formed *in vitro* (Bigalke *et al.*, 2014), in cells overexpressing PRV NEC (Hagen *et al*, 2015), and in perinuclear enveloped vesicles purified from HSV-infected cells (Newcomb *et al*, 2017). NEC oligomerization is necessary for budding because mutations intended to disrupt oligomeric interfaces reduce budding both *in vivo* and *in vitro* (Arii *et al*, 2019; Bigalke & Heldwein, 2015; Bigalke *et al.*, 2014; Roller *et al*, 2010). Collectively, these findings established the NEC as a robust membrane-budding machine that forms hexagonal scaffolds (reviewed in (Draganova *et al.*, 2020)).

Although NEC oligomerization is required for budding, NEC/membrane interactions may also have a mechanistic role in its budding mechanism. The TM helix of UL34 seemingly functions only to anchor the NEC to the INM (Schuster *et al*, 2012) because it is dispensable for budding *in vitro* (Bigalke *et al.*, 2014) and can be replaced with a heterologous TM *in vivo* (Schuster *et al.*, 2012). However, both UL31 and UL34 homologs have highly basic membrane-proximal regions (MPRs), and *in-vitro* budding by HSV-1 or PRV NEC requires acidic lipids (Bigalke *et al.*, 2014; Lorenz *et al*, 2015a), which implicates electrostatic interactions. Moreover, MPRs recruit the recombinant soluble HSV-1 NEC (which lacks the transmembrane (TM) anchor yet maintains robust budding activity) to acidic membranes *in vitro* (Bigalke *et al.*, 2014). It is yet unclear, however, how the MPRs interact with membranes or how these interactions lead to the formation of the negative membrane curvature during budding. Additionally, HSV-1 UL31 MPR is phosphorylated during infection (Chang & Roizman, 1993) by the viral kinase US3 (Kato *et al*, 2005) that targets six serines (Mou *et al*, 2009). The role of UL31 phosphorylation in nuclear egress is unclear, but phosphomimicking serine-to-glutamate mutations of these six serines inhibit nuclear egress and HSV-1 replication (Mou *et al.*, 2009), suggesting that phosphorylation may inhibit nuclear egress, by an unknown mechanism, presumably to prevent unproductive budding prior to the arrival of the capsid (reviewed in (Draganova *et al.*, 2020)). Thus, the MPRs may have both mechanistic and regulatory roles in NEC-mediated membrane budding. But it is unknown how the MPR/membrane interactions generate negative membrane curvature necessary for budding.

In addition to generating membrane buds, the NEC can also sever the necks of the budded vesicles at least in vitro (Bigalke *et al.*, 2014) and, potentially, in some infected cell types (Crump *et al*, 2007) even though in other cell types, the cellular ESCRT-III machinery is recruited for scission (Arii *et al*, 2018). Thus, another important unanswered question is how the NEC can generate both the membrane curvature necessary for the formation of the bud and a very different nanoscopic curvature required for scission to complete the budding process.

Here, by employing mutagenesis and several biophysical methods, we show that highly basic MPRs of the NEC are required for budding, can induce ordering within the headgroup and acyl chain regions of lipids in synthetic membranes, and can promote negative Gaussian curvature, which is the distinct type of curvature required for membrane scission. We propose that the NEC generates negative membrane curvature by a mechanism that combines lipid ordering and protein scaffolding. We also show that membrane remodeling by the NEC requires electrostatic interactions between the basic clusters within the MPRs and the acidic membranes. Further, we show evidence that the virus may control the membrane-budding activity of the NEC by manipulating its membrane interactions through phosphorylation, which would reduce the electrostatic interactions. Specifically, we demonstrate that the phosphomimicking mutations of serines adjacent to the basic clusters inhibit NEC-mediated budding in vitro, which explains how these mutations can also block capsid nuclear egress. HSV-1 may use phosphorylation to inhibit unproductive budding in the absence of the capsid by reducing the membrane-budding activity of the NEC.

## Results

### The MPRs are required for the efficient NEC-mediated membrane budding in vitro

HSV-1 UL31 is a soluble 306-amino-acid protein, and HSV-1 UL34 is a 275-amino-acid protein with a single C-terminal TM helix (Fig 1A). The highly basic MPRs encompass residues 1-50 of UL31 and 186-220 of UL34, which are absent from the crystal structures of the NEC cores and are located at the membrane-proximal end of the NEC (Fig 1B) (Bigalke & Heldwein, 2015). Previously, using an *in-vitro* budding assay with giant unilamellar vesicles (GUV) (Fig 1C), we showed that the NEC construct containing full-length UL31 and residues 1-220 of UL34 (NEC220) (Fig 1A) mediated robust membrane budding *in vitro* (Bigalke *et al.*, 2014). We also showed that the MPRs were necessary to recruit the NEC220 to synthetic membranes (Bigalke *et al.*, 2014) but did not investigate the potential role of the MPRs in the budding process beyond membrane recruitment partly because the soluble NEC220 must be recruited to the membranes from bulk solvent, making it difficult to uncouple NEC/membrane interactions necessary for budding from those necessary for membrane recruitment. To overcome this challenge, we utilized an NEC220 variant containing a C-terminal His_8_-tag in UL34 (Bigalke *et al.*, 2014). When used in conjunction with Ni-chelating lipids in the liposomes (Bubeck *et al*, 2005), polyhistidine tags efficiently tether proteins to membranes and are often used in place of TM anchors. The resulting NEC220-His_8_ construct had the same budding efficiency as the untagged NEC220 (Bigalke *et al.*, 2014).

**Fig. 1.**
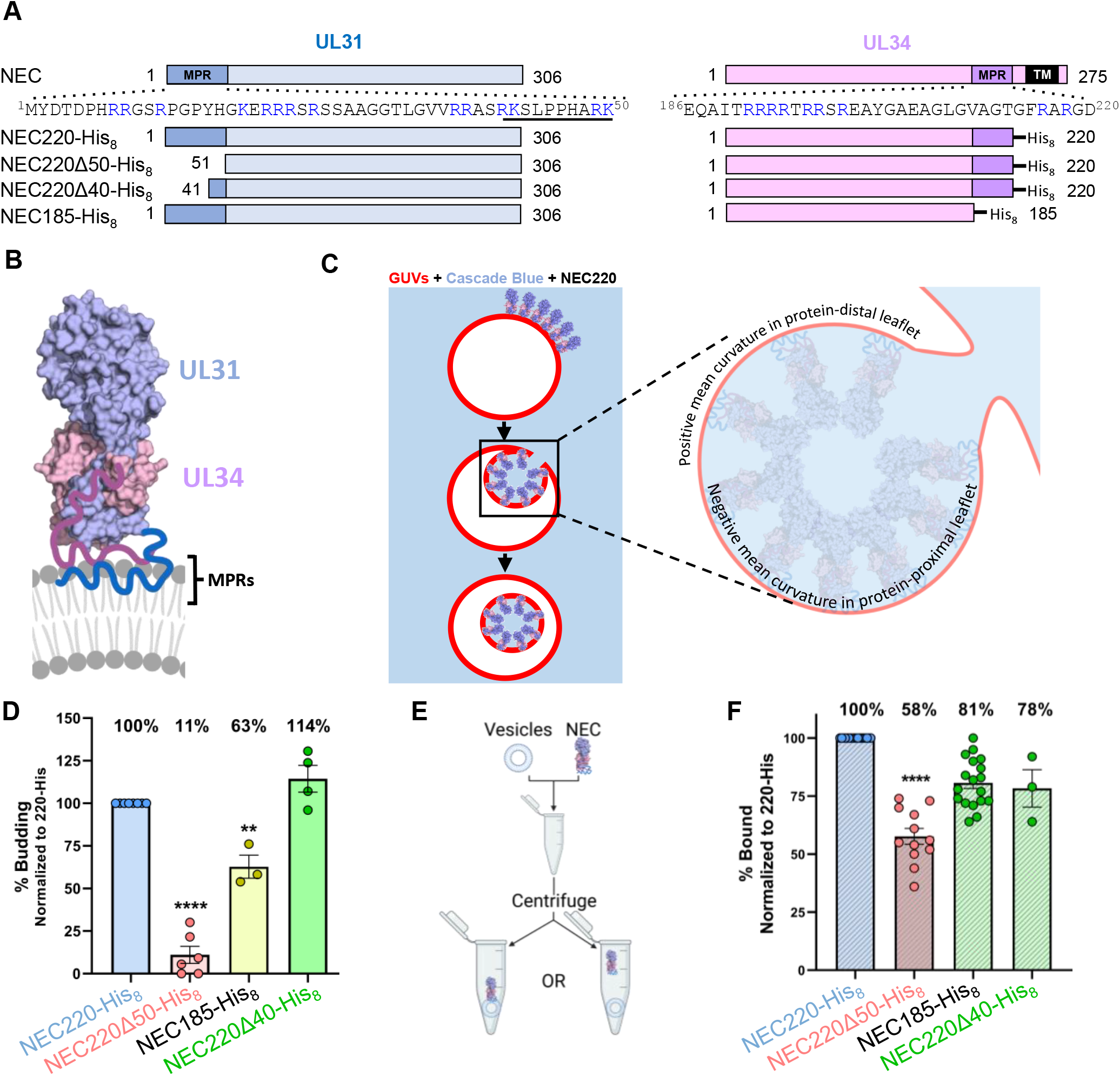
NEC MPRs are necessary for membrane vesiculation. (A) NEC construct map. Sequence of UL31 MPR residues 1-50 and UL34 MPR residues 186-220 shown at top. Basic residues in blue, UL31 mini-MPR is underlined. (B) Crystal structure of the NEC. MPRs missing from the structure and shown schematically in blue (UL31) and purple (UL34). UL34 TM is not included in schematic. Image generated using BioRender (BioRender.com). (C) *In vitro* budding assay. Red labeled GUVs are incubated with NEC in buffer containing Cascade Blue, a membrane-impermeant dye. Upon a budding event, intraluminal vesicles will form allowing blue dye into the red labeled GUV. Inset shows a budding vesicle depicting two types of mean curvature. Inset made with BioRender.com. (D) Vesicles contain Ni-chelating lipids to tether His_8_-tagged NEC to membranes. % budding was determined by counting the number of ILVs after addition of NEC and then normalized to NEC220-His_8_ amounts. Background levels of ILVs in the absence of NEC were subtracted from all values before normalization. Significance to 220∆40-His_8_ was calculated using an unpaired Student’s t-test with Welch’s correction (p<0.005=**, p<0.0005=***). In all plots, error bars represent the standard error of the mean (68% confidence interval of the mean) for at least two individual experiments. (E) *In vitro* co-sedimentation assay. Vesicles are incubated with NEC and then spun down in a centrifuge. Samples of the supernatant and pellet are run on an SDS-PAGE gel to determine the amount of NEC that pelleted with vesicles. Image made using Biorender.com. (F) % Bound was determined by quantification of SDS-PAGE gels of NEC +/− vesicles. Each bar represents the amount of protein pelleted. Significance to 220∆40-His_8_ was calculated using an unpaired Student’s t-test with Welch’s correction (p<0.0001=****).

By deleting the MPRs individually from the NEC220-His_8_ parent construct, we found that while both MPRs were required for efficient membrane budding *in vitro*, the UL31 MPR was more important because its deletion (NEC220Δ50-His_8_) reduced membrane budding to a very low level (11 ± 5% standard error of the mean, relative to NEC220-His_8_) whereas the deletion of the UL34 MPR (NEC185-His_8_) maintained budding at a moderate level (63 ± 7%) (Fig 1D). To assess the effect of MPR deletions on membrane recruitment, we used a co-sedimentation assay described previously (Fig 1E) (Bigalke *et al.*, 2014) with synthetic membranes lacking Ni-NTA-conjugated lipids but containing 40% negatively charged lipids, which are required for membrane recruitment of the soluble NEC220 (Bigalke *et al.*, 2014). NEC185-His_8_ associated with membranes more efficiently than NEC220Δ50-His_8_ (78 ± 8% vs. 58 ± 3% relative to NEC220-His_8_) (Fig 1F), which suggested that the UL31 MPR is more important for both membrane recruitment and budding activity than the UL34 MPR.

To narrow down residues within the UL31 MPR (Fig 1A) responsible for membrane interactions, we tested the truncation mutant NEC220Δ40-His_8_ that lacks residues 1-40 of the UL31 MPR (Fig 1A). Previously, we showed that these residues were dispensable for the membrane recruitment of soluble NEC220 (Bigalke *et al.*, 2014). Here, we found that these residues were also dispensable for budding (Fig 1D, F). Therefore, residues 41-50 can substitute for the full-length UL31 MPR during budding *in vitro*, and we refer to them as the “mini-MPR”.

### Basic clusters within the UL31 mini-MPR are essential for efficient budding

Due to its size, the mini-MPR of UL31 (^41^RKSLPPHARK^50^) provides an opportunity to dissect sequence requirements for NEC/membrane interactions and budding in a simplified system. Therefore, mutations were introduced into the NEC220Δ40-His_8_ parent construct. We first explored the role of the basic residues because electrostatic interactions between basic residues and acidic lipids commonly serve to recruit cytoplasmic proteins to membranes (Mulgrew-Nesbitt *et al*, 2006), and the MPRs of UL31 and UL34 homologs are rich in basic residues, 14 in HSV-1 UL31 (28%) and 9 in HSV-1 UL34 (31%) (Fig 1A and S1 Fig). Additionally, membrane binding by soluble HSV-1 NEC requires acidic lipids and is inhibited by high NaCl concentrations (Bigalke *et al.*, 2014), which further implicate electrostatic forces in NEC/membrane interactions.

The mini-MPR of UL31 has four basic residues arranged into two dibasic motifs, R41/K42 and R49/K50 (Fig. 2A), so we mutated them individually or together to serines, to maintain the polar character of the side chains (Fig 2B). Both dibasic motifs were required for efficient budding, with the first being more important than the second (Fig 2B) because the mutant containing only the first dibasic motif (NEC220Δ40-R49S/K50S-His_8_) maintained moderate budding efficiency (55 ± 10%) whereas the mutant containing only the second dibasic motif (NEC220Δ40-R41S/K42S-His_8_) budded as inefficiently (34 ± 10%) as the mutant lacking both dibasic motifs (NEC220Δ40-R41S/K42S/R49S/K50S-His_8_) (25 ± 6%) (Fig 2B). To probe the importance of charge distribution within the mini-MPR, we relocated the single dibasic motif, generating mutants NEC220Δ40-S43R/L44K-His_8_, NEC220Δ40-P45R/P46K-His_8_, and NEC220Δ40-H47R/A48K-His_8_. All three mutants mediated budding more efficiently than the mutants containing single dibasic motifs at either end of the mini-MPR (NEC220Δ40-R41S/K42S-His_8_ and NEC220Δ40-R49S/K50S-His_8_) (Fig 2B), which suggested that the location of the basic cluster can influence the budding efficiency.

**Fig. 2.**
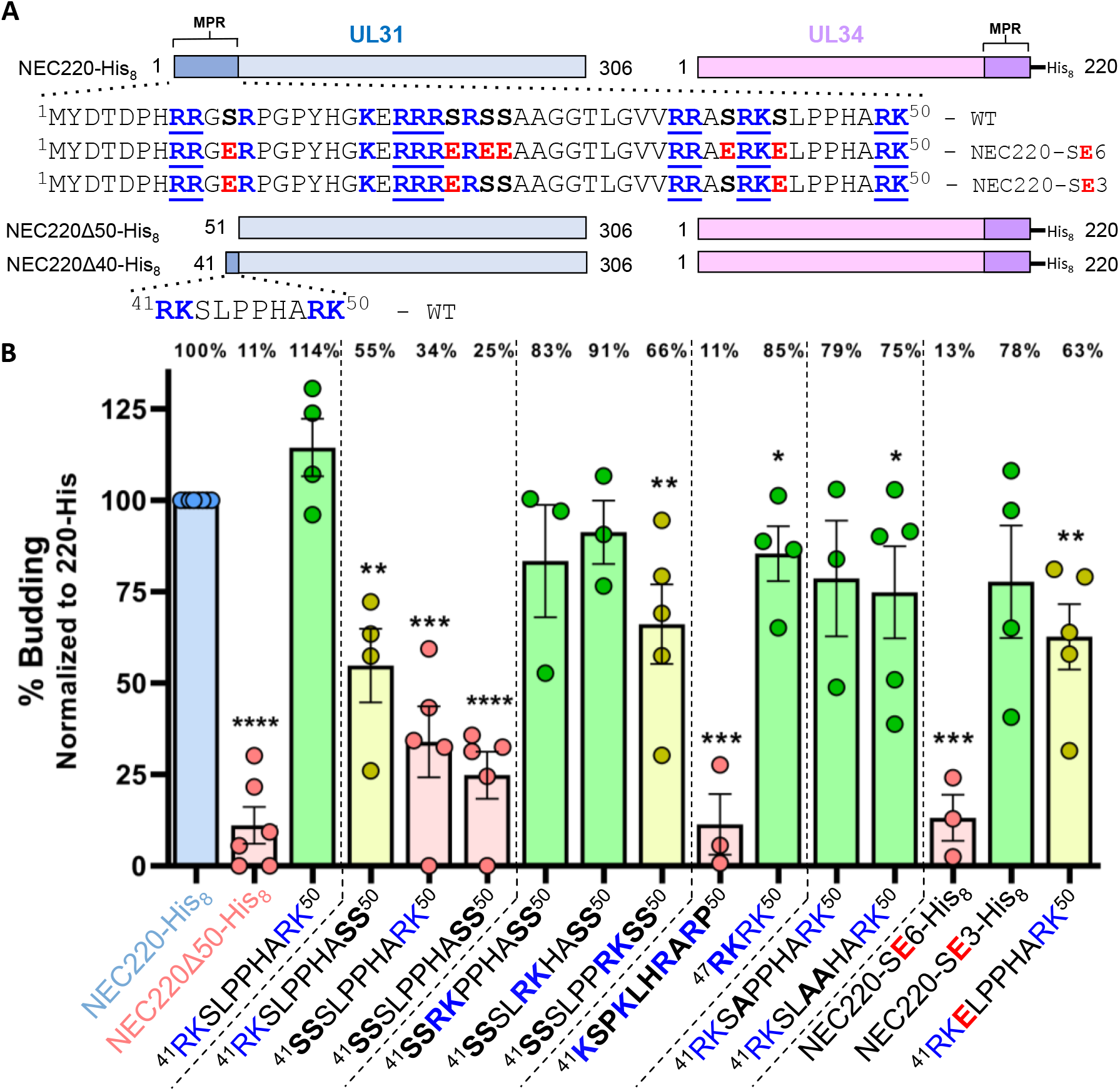
Altering the location of basic residues in the UL31 mini-MPR and introducing phosphomimicking mutations influence budding. (A) NEC construct map. Basic residues are shown in bold and blue, and clusters are underlined. Phosphorylatable serines are shown in bold. Glutamates in the phosphomimicking serine-to-glutamate mutants are shown in bold and red. (B) *In-vitro* budding assay. Mutated residues are shown in bold. % budding was determined by counting the number of ILVs after addition of NEC and then normalized to NEC220-His8 amounts. Background levels of ILVs in the absence of NEC were subtracted from all values before normalization. Significance to 220∆40-His8 was calculated using an unpaired Student’s t-test with Welch’s correction (p<0.05=*, p<0.005=**, p<0.0005=***, p<0.0001=****). In both plots, error bars represent the standard error of the mean (68% confidence interval of the mean) for at least three individual experiments. Coloring scheme based on significance: 0-49% is poor budding (red), 50-74% is moderate (yellow) and 75-100% is efficient (green). *In-vitro* co-sedimentation assay. % bound fraction was determined by quantification of SDS-PAGE gels of NEC +/− vesicles. Each bar represents the amount of protein pelleted. Binding values are to the right of the graph. Data is normalized to NEC220-, His_8_. Significance relative to NEC220∆40-His_8_ was calculated using an unpaired Student’s t-test with Welch’s correction (p<0.007=**, p<0.0001=****).

In the case of membrane association, a single dibasic motif sufficed for efficient membrane association (67 ± 15%, 80 ± 3%, and 81 ± 2%) unless it was located at the N terminus, in which case membrane association was similar to the mutant lacking basic residues (55 ± 11% and 50 ± 13%, respectively) (S2 Fig). Membrane association of NEC220Δ40-P45R/P46K-His_8_ could not be assessed because protein aggregated when incubated at room temperature for >15 minutes (S2 Fig). Distinct effects of dibasic motif mutations on budding vs. membrane association suggest that the requirements for efficient budding vs. membrane recruitment differ.

To probe the importance of charge clustering, we generated the scrambled mutant NEC220Δ40scr-His_8_ (^41^KSPKLHRARP^50^) that lacked basic clusters yet maintained the overall net +4 charge. This mutant associated efficiently with membranes (71 ± 5%) (S2 Fig) yet mediated budding at a minimal level (11 ± 8%) (Fig 2B) demonstrating the most pronounced difference between the requirements for budding vs. membrane recruitment. Thus, whereas membrane association requires a positive net charge of at least +2, membrane budding additionally requires charge clustering.

We also investigated the role of the LPP sequence in the middle of the mini-MPR. L44 is the sole hydrophobic residue within the mini-MPR, and hydrophobic interactions can contribute to protein/membrane interactions (Mulgrew-Nesbitt *et al.*, 2006), whereas the rigid di-proline motif in the middle of the mini-MPR could, in principle, adopt secondary structure important for membrane interactions. Yet, both the NEC220Δ40-L44A-His_8_ and the NEC220Δ40-P45A/P46A-His_8_ mutants supported efficient budding (Fig 2B), and therefore, the LPP sequence does not appear to play any role in either budding or membrane association.

To determine if 4 basic residues could replace the mini-MPR, we generated the NEC220Δ50-RKRK-His_8_ mutant. This mutant supported efficient budding (85 ± 8%) (Fig 2B) and membrane association (74 ± 8%) (S2 Fig). Thus, basic clusters are both necessary and sufficient for NEC-mediated budding *in vitro*. Similarly, the replacement of the UL31 MPR in PRV with 4 basic residues maintained efficient nuclear egress and replication of PRV (Klupp *et al*, 2018).

### Phosphomimicking mutations reduced both membrane association and budding

HSV-1 UL31 MPR is phosphorylated during infection (Chang &Roizman, 1993) by the viral kinase US3 (Kato *et al.*, 2005) that targets six serines, S11, S24, S26, S27, S40, and S43 (Mou *et al.*, 2009). The role of UL31 phosphorylation in nuclear egress has not yet been elucidated. Nevertheless, phosphomimicking mutations of these six serines (serine-to-glutamate) reduce nuclear egress and HSV-1 titers (Mou *et al.*, 2009), which suggests that phosphorylation may inhibit nuclear egress, by an unknown mechanism.

We have shown that positive charges in UL31 MPR are important for both the membrane association and the budding activity of the NEC. By decreasing the net positive charge of the UL31 MPR, the negative charges introduced by the phosphomimicking mutations would be expected to reduce both membrane association and the budding activity of the NEC. To test this, we generated the NEC220-SE6-His_8_ mutant, in which six serines within UL31 MPR were replaced with glutamates. Indeed, the phosphomimicking mutant had poor budding activity (13 ± 6%) (Fig 2B) and poor membrane association (23 ± 7%) (S2 Fig). To measure the effect of phosphomimicking mutations on budding in the context of the mini-MPR, which contains a single serine S43, we generated the NEC220Δ40-S43E-His_8_ mutant. The S43E mutation reduced budding (63 ± 9%) (Fig 2B) while preserving efficient membrane association (63 ± 8%) (S2 Fig), showing that adding a single negative charge to the UL31 mini-MPR impairs the budding ability of the NEC.

The location of basic clusters influences NEC membrane budding activity, so we hypothesized that the placement of phosphorylatable serines relative to basic residues may also be important for inhibition. Within the HSV-1 UL31 MPR, the 14 basic residues fall into five distinct clusters (Fig 2A), and each, except the C-terminal one, has at least one serine nearby (Fig 2A). To investigate whether single serine-to-glutamate mutations per cluster would recapitulate the inefficient budding phenotype of NEC220-SE6-His_8_, we generated the S11E/S24E/S43E mutant (NEC220-SE3-His_8_). However, NEC220-SE3-His_8_ supported efficient budding (78 ± 15%) (Fig 2B) and membrane association (70 ± 6%) (S2 Fig), showing that while adding six negative charges was sufficient to inhibit budding, adding three was not. The budding ability of the NEC thus requires not only basic clusters but also a sufficiently high net positive charge within the UL31 MPR.

Collectively, these results show that phosphomimicking mutations within the UL31 MPR, which introduce negative charges, reduce its budding activity, which confirms the importance of the net positive charge within the UL31 MPR for the NEC function. We propose that the impaired nuclear egress and reduced titers of the phosphomimicking HSV-1 NEC mutant *in vivo* (Mou *et al.*, 2009) is due to its reduced budding activity. Phosphorylation, which also introduces negative charges, would be expected to have a similar inhibitory effect on budding. We hypothesize that by inhibiting the budding activity of the NEC, phosphorylation could serve to prevent unproductive budding prior to the arrival of the capsid (reviewed in (Draganova *et al.*, 2020)).

### Soluble NEC inserts peripherally into the tethered lipid bilayers

To determine the orientation of the NEC on the membrane and how deeply it inserts into the lipid bilayer, we turned to neutron reflectometry (NR) (Vanegas *et al*, 2018), which allows low-resolution structural characterization of the membrane and any associated protein. A tethered lipid bilayer composed of POPC/POPS/POPA in a 3/1/1 molar ratio was prepared in a flow cell, and the reflectivity of the bilayer interface to a collimated neutron beam, incident at various angles, was measured before and after incubation with NEC220 at 100 nM or 500 nM (S3 Fig). Protein density profiles calculated from the NR measurements at each NEC concentration overlapped only with the density profile of the outer lipid headgroup (Fig 3B), suggesting that NEC220 inserted only into the polar lipid headgroup region (Fig 3A and B), without penetration of large domains into the acyl chain region.

**Fig. 3.**
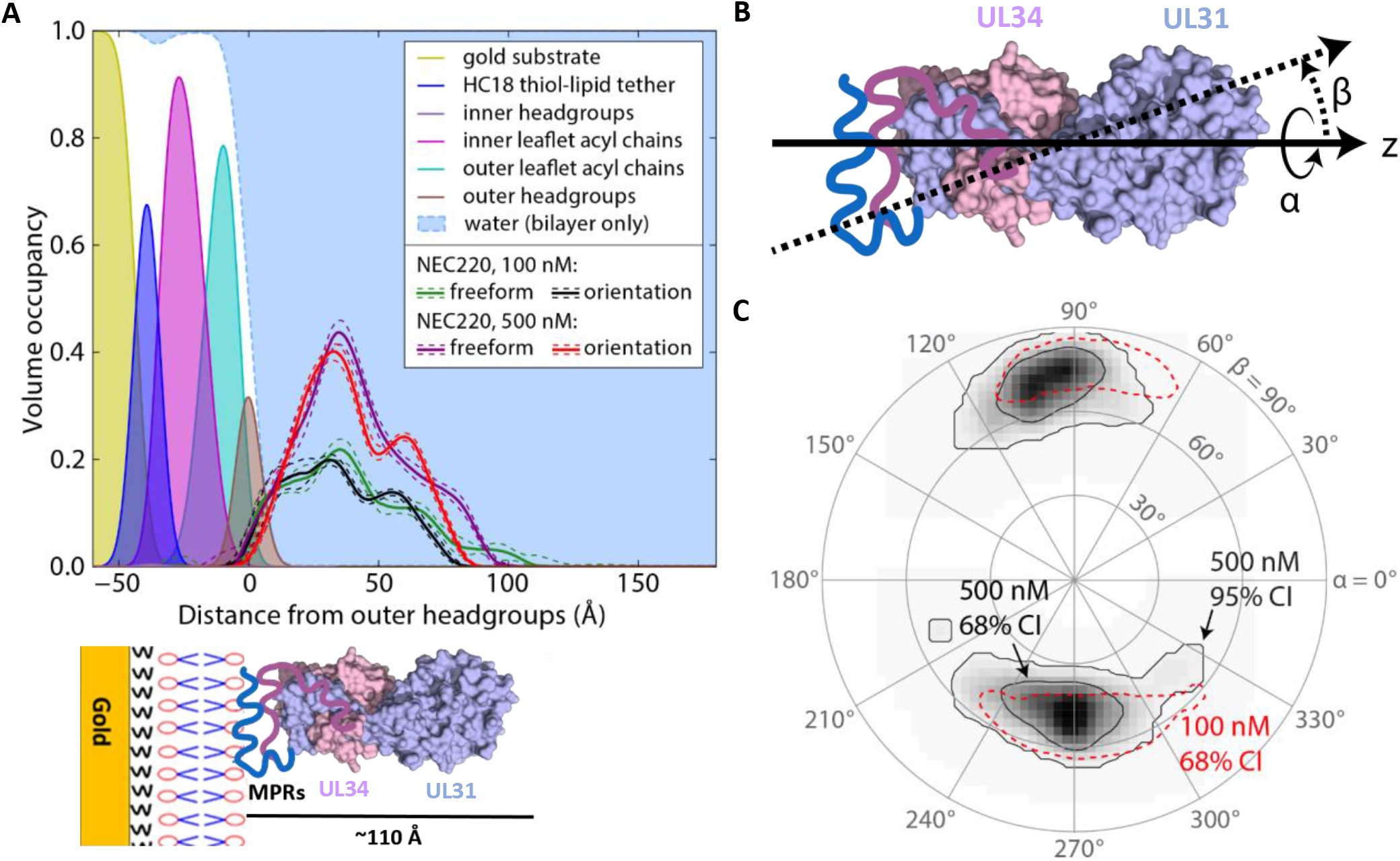
NEC inserts into polar lipid headgroups. (A) Volume density profiles of 3:1:1 mol% POPC:PS:PA lipid membranes determined from fitting a composition space model to the NR spectra. Density profiles of substrate and bilayer components are shown by filled curves; the sum is shown by the dashed blue line, and water fills the remaining space. Protein density profiles derived from freeform (Catmull-Rom spline) and orientation (Euler rotations of the crystallographic structure) models after incubation with 100nM and 500nM bulk concentrations of NEC220 and subsequent buffer rinses. Dashed lines indicate 68% confidence intervals on the protein density profiles. Schematic underneath graph is shown to provide context for each peak in graph. (B) Euler angle rotation scheme. (C) Probability plot for the orientation of NEC220 at the membrane as parameterized by the Euler angles α and β shown in (B). The contour lines represent the 68% and 95% confidence intervals, as labeled.

Within the NEC coats formed *in vitro* and *in vivo*, the NEC molecules are oriented perpendicularly to the plane of the membrane, with the protein density extending ~110 Å from the membrane, in accordance with the cryo-EM measurements (Bigalke *et al.*, 2014; Hagen *et al.*, 2015). However, the NEC220 density profile obtained from the NR measurements only extended to ~90 Å, and an orientation probability plot showed significant tilt of the NEC220 from a vertical orientation (Fig 3C). These data seemingly suggest that on the NR substrates, the NEC220 adopts a tilted orientation relative to the plane of the membrane. However, because NR data are averaged over both time and space, they likely reflect different states of the NEC characterized by different levels of positional freedom, for example, individual heterodimers vs. higher oligomers. We hypothesize that whereas the individual NEC heterodimers can adopt a range of orientations relative to the plane of the membrane, oligomerization into hexagonal patches, or even individual hexamers, would restrict the movement of the NEC molecules resulting in a more upright NEC density profile. We note that the intrinsic flatness of the NR substrates, or alternatively the underlying grain structure of the gold, may prevent the formation of extended hexagonal coats.

We also observed that after exposure to 500 nM NEC220, which deposited NEC220 on the membrane surface at high density (molar ratio protein/lipid (P/L) = 1/45), the membrane thickened by 0.49 ± 0.17 Å (68% confidence interval), in the context of the orientation model (Table S1). Thinning of membranes tethered to flat substrates has been observed with proteins that generate positive curvature by inserting into the headgroup region (Chen *et al*, 2003; Mihailescu *et al*, 2014; Mihailescu *et al*, 2019). This is because forcing the headgroups apart on a flat substrate increases the area per lipid and thins the membrane (the hydrophobic tails form a constant-volume cylinder, the height of which must decrease if the area is increased).

Conversely, membrane thickening could occur if the headgroups were forced closer together on a flat substrate, which on free membranes would result in negative mean membrane curvature. We hypothesize that the ability of the NEC to generate negative membrane curvature manifests as membrane thickening on the NR substrates.

### NEC UL31 MPR peptides induce lipid headgroup ordering

To determine how the MPRs influence the structure of the lipid bilayer, we turned to continuous-wave electron spin resonance (CW-ESR) using spin-labeled lipids, which generate an ESR signal. The spin-labeled lipid within the membrane is sensitive to the local environment, and, therefore, the ESR signal reports on the mobility of the spin label, which, in turn, reports on the order of the lipids in the membranes. The order parameter of the spin (S_0_), which is calculated from the CW-ESR spectra, correlates with the local lipid order and inversely correlates with the mobility of the spin label. Thus, the effect of peptide binding on the lipid order can be monitored. Two phosphatidylcholine derivatives containing spin labels were used: dipalmitoylphosphatidyl-tempo-choline (DPPTC), which has a tempo-choline headgroup with a spin sensitive to the environment within the headgroup region (Fig 4B), and 1-palmitoyl-2-stearoyl-(5-doxyl)-sn-glycero-3-phosphocholine, which has a doxyl group in the C5 position of the acyl chain where the spin is sensitive to the environment within the upper acyl chain (Fig 4C). These two spin-labeled lipids were used in previous studies, which validated their ability to detect changes in lipid order (Ge & Freed, 2003, 2009, 2011; Lai & Freed, 2014; Lai & Freed, 2015; Lai *et al*, 2017; Nathan *et al*, 2020; Pinello *et al*, 2017; Straus *et al*, 2019).

**Fig. 4.**
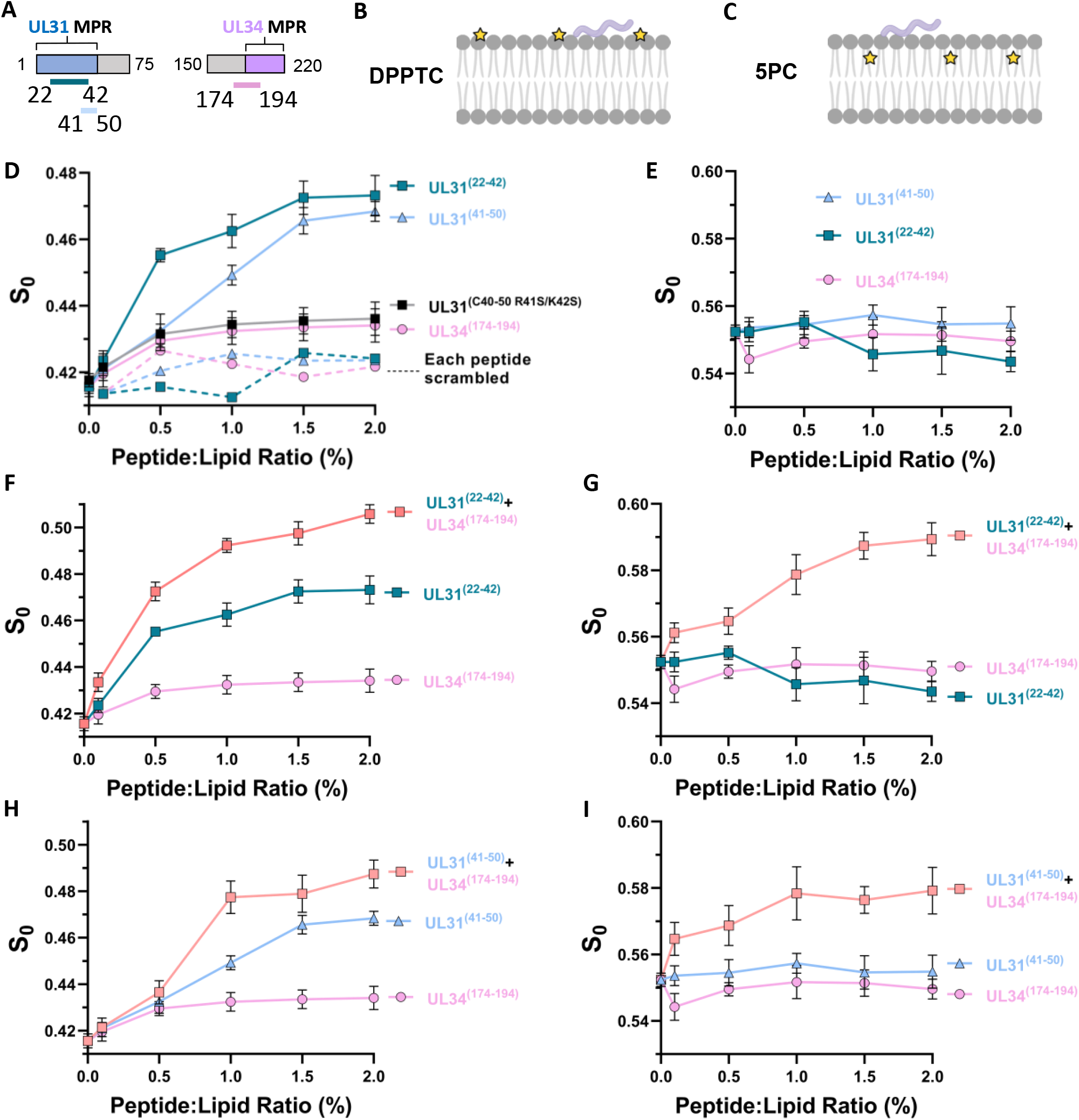
UL31 and UL34 membrane proximal region peptides induce lipid headgroup and acyl chain ordering. (A) Schematic depicting peptide location within the NEC sequence. (B) Schematic depicting DPPTC (yellow star is the probe) spin-labeled lipid in membrane and peptide (light purple). Image created with BioRender.com. (C) Schematic depicting 5PC spin-labeled lipid in membrane with peptide. Plot of order parameters of DPPTC in POPC/POPS/POPA 3:1:1 versus the P/L ratio of individual UL31^(22-42)^ (teal), UL31^(41-50)^ (light blue), UL34^(174-194)^ (light pink) and UL31R41S/K42S^(41-50)^ (black). Scrambled peptides shown with dashed lines, color coded as previously stated. In all plots, error bars represent standard deviation (68% confidence interval of the data) of at least three individual experiments. (E) Plot of order parameters of 5PC in POPC/POPS/POPA 3:1:1 versus the P/L ratio of individual UL31^(22-42)^, UL31^(41-50)^, UL34^(174-194)^. (F) Plot of order parameters of DPPTC in POPC/POPS/POPA 3:1:1 versus the P/L ratio of individual UL31^(22-42)^, UL34^(174-194)^ and combination of the two (light orange). (G) plot of order parameters of 5PC in POPC/POPS/POPA 3:1:1 versus the P/L ratio of UL31^(22-42)^, UL34^(174-194)^ and combination of the two. (H) plot of order parameters of DPPTC in POPC/POPS/POPA 3:1:1 versus the P/L ratio of individual UL31^(41-50)^, UL34^(174-194)^ and combination of the two. (I) Plot of order parameters of 5PC in POPC/POPS/POPA 3:1:1 versus the P/L ratio of individual UL31^(41-50)^, UL34^(174-194)^ and combination of the two.

To investigate the effect of the NEC MPRs on lipid order, we used three UL31-derived peptides: UL31^(41–50)^, which corresponds to the mini-MPR; UL31^(C40-50^ ^R41S/K42S)^, which corresponds to the mini-MPR with the mutated N-terminal dibasic motif and contains an N-terminal cysteine for spin-labeling in later experiments (Fig 2); and UL31^(22–42)^, which corresponds to the middle of the UL31 MPR. We also used one UL34-derived peptide, UL34^(174–194)^, which encompasses a portion of the UL34 MPR (Fig 4A). The boundaries of UL31^(22–42)^ and UL34^(174–194)^ were chosen using a machine-learning classifier that identifies peptide sequences with the capacity to generate negative Gaussian curvature in membranes, which is topologically required in membrane-remodeling processes such as membrane budding and fission (Lee *et al*, 2016). As controls, we also prepared scrambled versions of the peptides: UL31^scr(41-C51)^, UL31^scr(22-C43)^, and UL34^scr(174–194)^. Scrambled UL31 peptides contained C-terminal cysteines for spin-labeling in later experiments. Peptide sequences are listed in S2 Table.

If peptide binding to the membrane increases the mobility of the spin-labeled probe, we would expect to see a decrease in the order parameter, S_0_, with increasing peptide/lipid (P/L) ratio. Conversely, if peptide binding decreases the mobility of the spin-labeled probe, we would see an increase in S_0_ (Ge & Freed, 2003). All three native peptides UL31^(41–50)^, UL31^(22–42)^, and UL34^(174–194)^ increased the S_0_ in the headgroup region (DPPTC) (Fig 4D) in a sequence-specific manner, with the native UL31 peptides inducing significantly larger lipid headgroup ordering than the scrambled versions. However, none of the individual MPR peptides induced obvious ordering of the upper acyl chain (5PC) (Fig 4E). The CW-ESR experiments also showed that the UL31^(C40-50 R41S/K42S)^ mutant peptide, which lacks the N-terminal dibasic motif, induced substantially less lipid headgroup ordering than the WT UL31^(41–50)^ peptide (Fig 4D).

Decreased lipid headgroup ordering by the UL31^(C40-50 R41S/K42S)^ and the UL31^scr(41-C51)^ peptides (Fig 4D) correlates with the reduced budding activity of the respective mutant NEC constructs NEC220Δ40-R41S/K42S-His_8_ (34 ± 10%) and NEC220Δ40scr-His_8_ (11 ± 8%) (Fig 2B). Decreased lipid headgroup ordering by the UL31^(C40-50 R41S/K42S)^ mutant peptide could potentially be due to reduced membrane binding (relative to UL31^(41–50)^) as determined by the ESR partition ratio (S4 Fig). However, the UL31^scr(41-C51)^ peptide binds membranes similarly to UL31^(41–50)^ (S4 Fig), so the observed decrease in lipid headgroup ordering could not be due to impaired membrane interactions. These results suggest that both lipid ordering (Fig 4D) and efficient budding *in vitro* (Fig 2B) require not only the +4-net charge, but charge clusters, namely, 2 dibasic motifs.

### In combination, UL31 and UL34 MPR peptides induce both lipid headgroup and acyl chain ordering

We next examined how a combination of UL31 and UL34 MPR peptides would affect lipid order. A mixture of UL31 and UL34 MPR peptides at a 1/1 molar ratio was mixed with liposomes containing spin-labeled lipids in various P/L ratios. When comparing S_0_ at the same P/L ratio, the UL31^(22–42)^/UL34^(174–194)^ combination increased the local order in the headgroup region (DPPTC) to a greater extent than the individual peptides alone (Fig 4D and F). The same effect was observed for the UL31^(41–50)^/UL34^(174–194)^ combination (Fig 4D and H). Moreover, both the UL31^(22–42)^/UL34^(174–194)^ and the UL31^(41–50)^/UL34^(174–194)^ combinations induced ordering of the upper acyl chains (5PC, Fig 4G and I), in contrast to the individual peptides. Therefore, while individually, UL31 and UL34 MPR peptides induce lipid headgroup ordering, in combination, they induce greater lipid headgroup ordering as well as the ordering of the upper acyl chains. Thus, the UL31 and UL34 MPR peptides act cooperatively.

The ESR measurements were also performed with NEC220 and NEC220Δ40. Both protein complexes induced membrane ordering in the headgroup region, with NEC220 having a larger effect than the NEC220Δ40 (S5 Fig). The “nominal” P/L ratio of the complex required to saturate the S_0_-P/L ratio curve was significantly larger than that of the peptide mixtures, which could be due to the different binding constants of the peptides relative to the NEC constructs. The ESR experiments utilized small unilamellar vesicles (SUVs), <100 nm in diameter, which both NEC220-His_8_ and NEC220Δ40-His_8_ bind less efficiently than lipid vesicles of larger size (S5 Fig). Thus, the CW-ESR results show that both the MPR-derived peptides and the NEC can induce membrane ordering.

### In the presence of the UL34 MPR, the UL31 MPR inserts more deeply into the membrane

To measure how deeply the MPR peptides insert into the membrane, we performed power saturation ESR (Georgieva *et al*, 2010; Georgieva *et al*, 2014; Snead *et al*, 2017) with peptides spin-labeled with S-(1-oxyl-2,2,5,5-tetramethyl-2,5-dihydro-1H-pyrrol-3-yl) (MTSL) on either an N-terminal cysteine (UL31^(C40-50)^ and UL31^(C21-42)^) or a C-terminal cysteine (UL31^(41-C51)^ and UL31^(22-C43)^). The depth of spin label insertion into the membrane was determined from the accessibility of each peptide to O_2_, which penetrates into the hydrophobic region of the membrane, or Ni(II)‐diammine‐2,2’‐(ethane‐1,2‐diyldiimino) diacetic acid (NiEDDA), which does not penetrate the membrane beyond the polar headgroup region. The insertion depth parameter Φ, which represents the difference in the accessibility of the spin label to O_2_ vs. NiEDDA, reports on the spin label insertion depth, with Φ = 0 corresponding to the hydrophobic/hydrophilic interface. Thus, the more positive the Φ, the deeper the residue inserts into the hydrophobic core, whereas a negative Φ means the residue remains in the polar headgroup region.

The Φ values for the spin-labeled UL31^(C40-50)^ and UL31^(41-C51)^ were −0.41 ± 0.03 (68% confidence interval) and −0.69 ± 0.04, respectively (Fig 5), which indicated that they both reside in the lipid headgroup region. However, when the spin-labeled UL31^(C40-50)^ and UL31^(41-C51)^ peptides were mixed with the unlabeled UL34^(174–194)^ peptide in a 1:1 molar ratio, the Φ values increased to −0.09 ± 0.04 and −0.18 ± 0.04, respectively, consistent with a deeper insertion into the hydrophobic/hydrophilic interface (Fig 5). A similar trend was observed for UL31^(C21-42)^ and UL31^(22-C43)^, where the Φ values of the spin-labeled cysteines increased from −0.30 ± 0.03 and −0.42 ± 0.03 to 0.17 ± 0.04 and 0.12 ± 0.02, respectively, in the presence of the unlabeled UL34^(174–194)^ peptide (Fig 5). The power saturation ESR results suggest that the UL31 MPR inserts more deeply into the membrane in the presence of UL34 MPR. This observation complements the CW-ESR results, which showed that the 1:1 mix of UL31 and UL34 MPR peptides induces lipid ordering within the upper acyl chains (Fig 4), which could result from the deeper insertion of the UL31 MPR into the upper acyl chain region in the presence of the UL34 MPR. Alternatively, the UL31 MPR may remain in the lipid headgroup region while drawing the headgroups together and thereby constraining the motion of the upper acyl chains and the spin label located there.

**Fig. 5.**
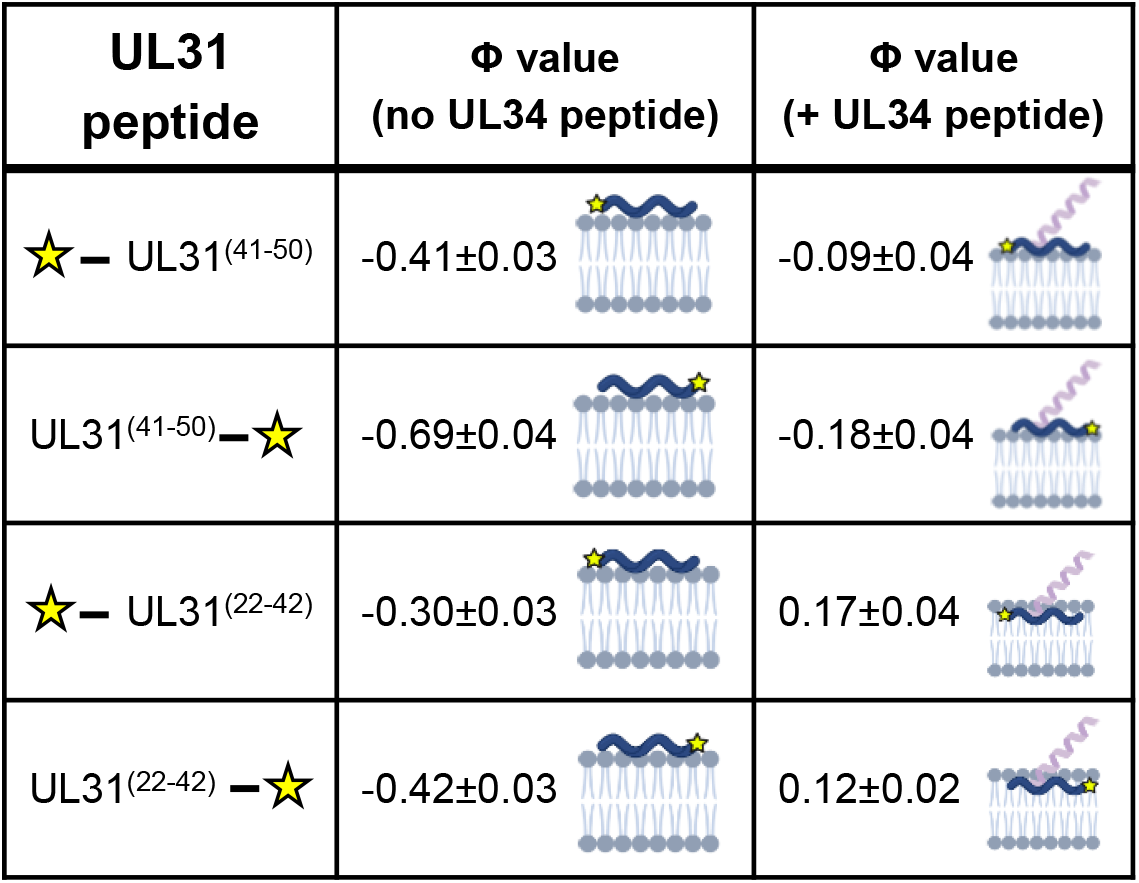
Presence of the UL34 peptide deepens membrane penetration of UL31 MPR peptides. The insertion depth parameter Phi (Φ) values of the N- and C-terminal spin labeled UL31 MPRs in the presence of POPC/POPS/POPA=3/1/1 SUVs. The Phi values were calculated from the power saturation ESR spectra. The averages and standard deviations (68% confidence intervals of the data) were calculated from three independent experiments. Schematics to the right of Φ values depict probe (yellow star) placement on peptides and roughly estimated insertion depths. Images created with BioRender.com.

### UL31 and UL34 MPRs induce negative Gaussian curvature in membranes

To determine the effect of the MPRs on membrane curvature, we used small-angle X-ray scattering (SAXS) to quantitatively characterize membrane deformations upon exposure to MPR-derived peptides UL31^(22–42)^, UL31^(41–50)^, UL34^(174–194)^, and their combinations. SAXS can detect the generation of negative Gaussian curvature (NGC) (Kaplan *et al*, 2017; Mishra *et al*, 2011; Schmidt *et al*, 2011; Schmidt *et al*, 2013b), which corresponds to the saddle-like curvature found on the inside of a donut hole, the inner surface of membrane pores, and the necks of budding vesicles (Fig 6A) and is required for membrane-remodeling events such as vesicle budding (Schmidt *et al.*, 2013b), membrane fission (Lee *et al*, 2017), membrane fusion (Yao *et al*, 2015), and pore formation (Schmidt *et al.*, 2011). By contrast, positive Gaussian curvature (PGC) corresponds to the dome-like curvature such as found on a spherical body of the bud (Fig 6A).

**Fig. 6.**
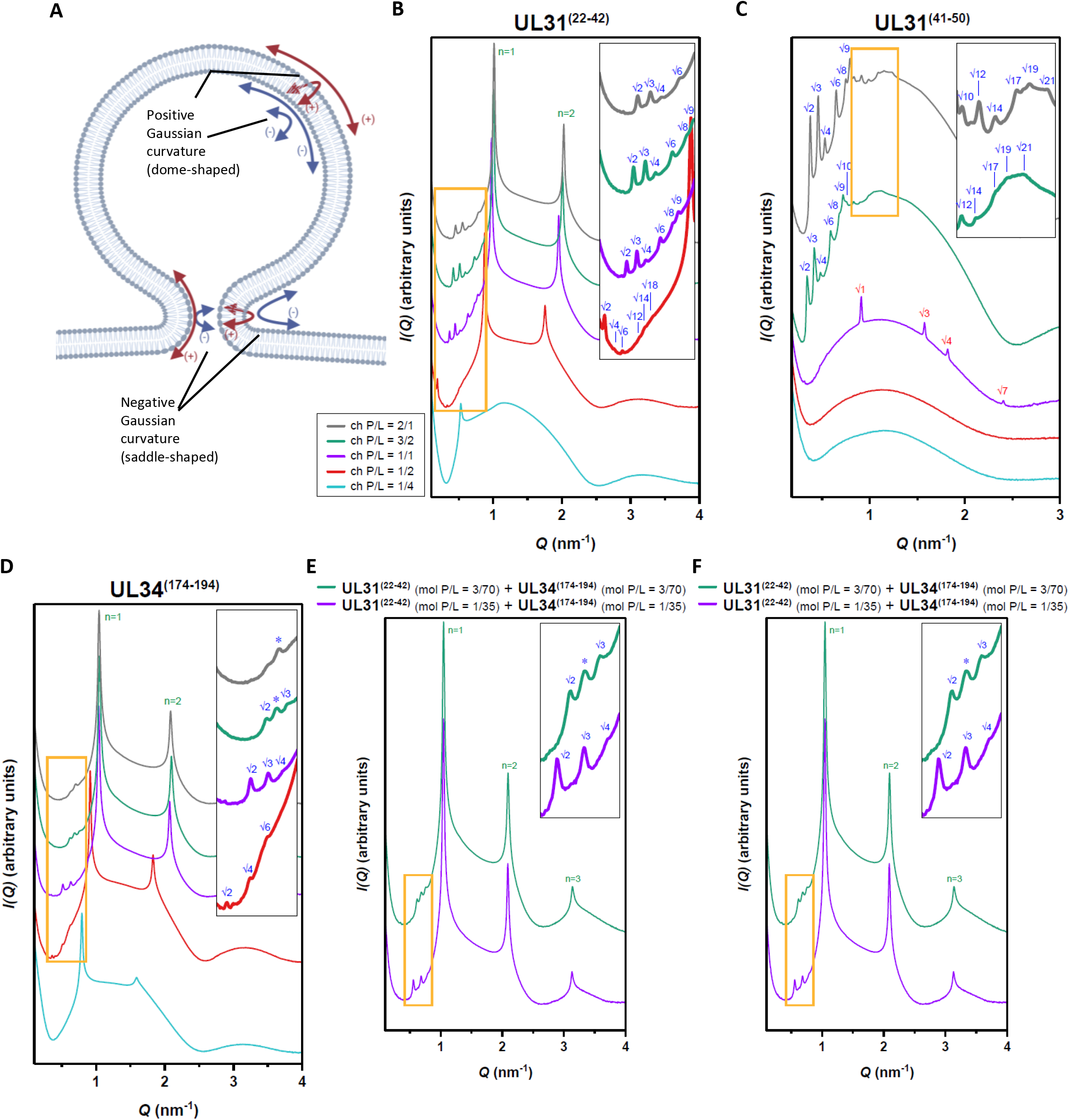
UL31 and UL34 membrane proximal region peptides generate negative Gaussian curvature in membranes. SAXS spectra for SUVs incubated with individual peptides (A) Schematic depicting the principal curvatures of the neck of a budding vesicle which together generate negative Gaussian curvature. Image created with BioRender.com. (B) UL31^(22-42)^, (C) UL31^(41-50)^, (D) UL34^(174-194)^, and two combinations of approximately equimolar amounts of each UL31 and UL34 MPR peptide (E) UL31^(22-42)^/ UL34^(174-194)^, and (F) UL31^(41-50)^/ UL34^(174-194)^. (A–F) For improved visualization, spectra have been manually offset in the vertical direction by scaling each trace by a multiplicative factor. For clarity, the insets show expanded views of the lower intensity cubic reflections (orange-boxed regions). Indexed reflections for cubic (blue), hexagonal (red), and lamellar (green) phases are labeled. Asterisks denote peaks that could not be indexed to a phase due to absence of higher order reflections.

SUVs with a 1/4 molar ratio of DOPS/DOPE were incubated with each peptide or combination of peptides at peptide-to-lipid charge ratios (ch P/L) of 1/4, 1/2, 1/1, 3/2, and 2/1 (see Methods for the equivalent peptide-to-lipid molar ratios [mol P/L]) and measured using SAXS. We choose a lipid composition DOPS/DOPE = 20/80 because it has a surface charge density typical of eukaryotic membranes and can sense the capacity for the induction of membrane curvature, including NGC. The induction of NGC was monitored by the appearance of correlation peaks that correspond to NGC-rich *Im3m* and *Pn3m* cubic phases, which are defined by a lattice parameter *a* and an average NGC |<*K*>| (Fig 6, S7 Fig). Both *Im3m* and *Pn3m* are inverse bi-continuous cubic phases (Q_II_), which are lyotropic liquid-crystalline phases that can be formed by lipid systems. A bi-continuous cubic phase consists of two interpenetrating, but non-intersecting, aqueous volumes that are separated by a single continuous lipid bilayer. The mid-plane of this bilayer traces a minimal surface that is characterized by having NGC at all points on its surface.

While the SUVs alone displayed a broad characteristic feature consistent with the form factor expected for unilamellar vesicles (S6 Fig), all three individual peptides and the two peptide mixtures restructured the membranes into NGC-rich cubic phases with different amounts of NGC (Fig 6B–D, S7 and S8 Figs.). The NGC magnitude generally increased with increasing peptide concentration. Among the three individual peptides, UL34^(174–194)^ induced the highest amounts of NGC on average, followed by UL31^(22–42)^ and UL31^(41–50)^. While all three individual peptides were able to form cubic phases, over five times the number of UL31^(41–50)^ peptide molecules were required to generate approximately the same quantitative amount of NGC as UL31^(22–42)^ or UL34^(174–194)^, which suggests that UL31^(41–50)^ peptide has a reduced capacity for NGC induction compared with the other two peptides.

Upon exposure to these peptides, in addition to the cubic phases, the membranes tended to form additional coexisting phases, which suggested the presence of other modes of membrane deformation. Interestingly, at ch P/L = 1/1, UL31^(41–50)^ formed an inverse hexagonal phase (H_II_), which is characterized by having negative mean curvature (but zero Gaussian curvature). This property is in line with the requirement of the UL31 MPR for budding (Fig. 1D). Additionally, both UL31^(22–42)^ and UL34^(174–194)^ but not UL31^(41–50)^ induced co-existing lamellar phases (L_μ_) (Fig 6A, S8 Fig), but the relevance of these to the topological changes that occur during budding, if any, is unclear.

We further examined the membrane curvature effects of peptide combinations, UL31^(22–42)^/UL34^(174–194)^ and UL31^(41–50)^/UL34^(174–194)^. At approximately equimolar ratios, both peptide pairs generated higher magnitudes of NGC than the individual peptides (Fig 6EF, S7DE, S8 Fig), demonstrating a cooperative effect between the UL31 and UL34 MPR peptides, which is consistent with their cooperativity in inducing lipid ordering observed by the ESR. This is also consistent with previous studies that showed that embedded peptides and proteins introduce intramembrane stresses and strains that lead to negative curvature generation and alter membrane bending stiffness (Campelo *et al*, 2008a; Zemel *et al*, 2008). Thus, while the UL31 and UL34 MPR peptides can generate NGC as individual peptides, they do so more effectively when combined. Using a catenoid surface model (Kozlovsky & Kozlov, 2003; Lee *et al.*, 2017; Schmidt *et al.*, 2013b), we estimated the size of the constricted membrane neck of a budding vesicle that can be formed from the highest amount of NGC induced by the UL31 and UL34 MPR peptides to be |<*K*>| = 3.21 × 10^-2^ nm^-2^, which corresponds to a membrane neck with an inner diameter of 7.2 nm and an outer diameter of 15.2 nm (assuming ~4 nm thick membrane bilayer). This estimate is in agreement with the diameters of the scission necks formed by mitochondrial fission proteins (Lee *et al.*, 2017) and with the theoretical calculations (Kozlovsky &Kozlov, 2003). Thus, the MPRs can generate membrane curvature necessary for neck scission, which is consistent with the NEC-induced bud scission observed *in vitro* (Bigalke *et al.*, 2014).

### DEER analysis suggests that UL31 and UL34 MPRs interact on membranes

The cooperative effect of the UL31 and UL34 MPR peptides on lipid ordering and induction of NGC as well as a greater depth of insertion of the UL31 MPR peptides in the presence of UL34 MPR peptide suggest that the UL31 and the UL34 MPR peptides interact. To determine if the UL31 and UL34 peptides interact on membranes, we employed double electron-electron resonance (DEER) spectroscopy, which yields the distance distributions between two spin systems in a frozen sample and is sensitive within the 20-80 Å range (Borbat & Freed, 2007; Borbat *et al*, 2013). The recently developed pulse-dipolar electron-spin resonance spectroscopy wavelet denoising methodology removes the noise from the ESR spectra and improves their accuracy (Srivastava *et al*, 2017) thereby reducing the uncertainty in distance distribution reconstruction by a special singular value decomposition methodology (Srivastava & Freed, 2017, 2019).

In our experiments, one spin was attached to either the N-or the C-terminal cysteine of a UL31 peptide and the other, to the native cysteine, C182, of the UL34 MPR peptide. None of the individual peptides exhibited any DEER signal in the presence of SUVs in 1/200 P/L ratio (see representative DEER spectra of UL31^(C1-50)^ and UL34^(174–194)^ in Figure S9AB). In a spin echo control experiment, strong spin echoes were observed (S9C Fig), which indicated that the peptides were properly spin-labeled and did not aggregate, ruling out the possibility that the phase memory time (T_m_) were too short to observe the DEER signal. Therefore, a lack of a DEER signal with individual peptides indicates that they do not homodimerize in the presence of SUVs (Georgieva *et al*, 2015).

Next, we mixed each of the six singly labeled UL31 MPR peptides (UL31^(C1-50)^, UL31^(1-C51)^, UL31^(C40-50)^, UL31^(41-C51)^, UL31^(C21-42)^, or UL31^(22-C43)^) with the singly labeled UL34 MPR-peptide in 1/1 ratio in the presence of SUVs. DEER measurements between UL34^(174–194)^ and UL31^(41-C51)^ or UL31^(1-C51)^ were similar, 23.0 ± 0.17Å (68% confidence interval of the mean) and 27.6 ± 0.16 Å, respectively (Fig 7E), which suggests that the mini-MPR recapitulates the interactions of the full-length UL31 MPR. Additionally, residues C40_31_ and C51_31_ are equidistant from C182_34_ (23.6 ± 0.13 Å and 23.0 ± 0.17/27.6 ± 0.16 Å, respectively) whereas both C21_31_ and C0_31_ are much farther away (42.9 ± 0.08 Å and 49.7 ± 0.20 Å, respectively) (Fig 7E). This suggests that the UL31 MPR C terminus is closer to the UL34 MPR than its N terminus and likely interacts with it. The C43_31_-C182_34_ distance (35.5 ± 0.06 Å) is unexpectedly longer than both the C40_31_-C182_34_ and the C51_31_-C182_34_ distances (23.6 ± 0.13 Å and 23.0 ± 0.17/27.6 ± 0.16 Å, respectively) (Fig 7E), but this could be due to the relative orientations of the spins, which are ~6-Å away from the Cμ (Alexander *et al*, 2013). As a control, no DEER signal was observed in the absence of SUVs (S9D).

**Fig. 7.**
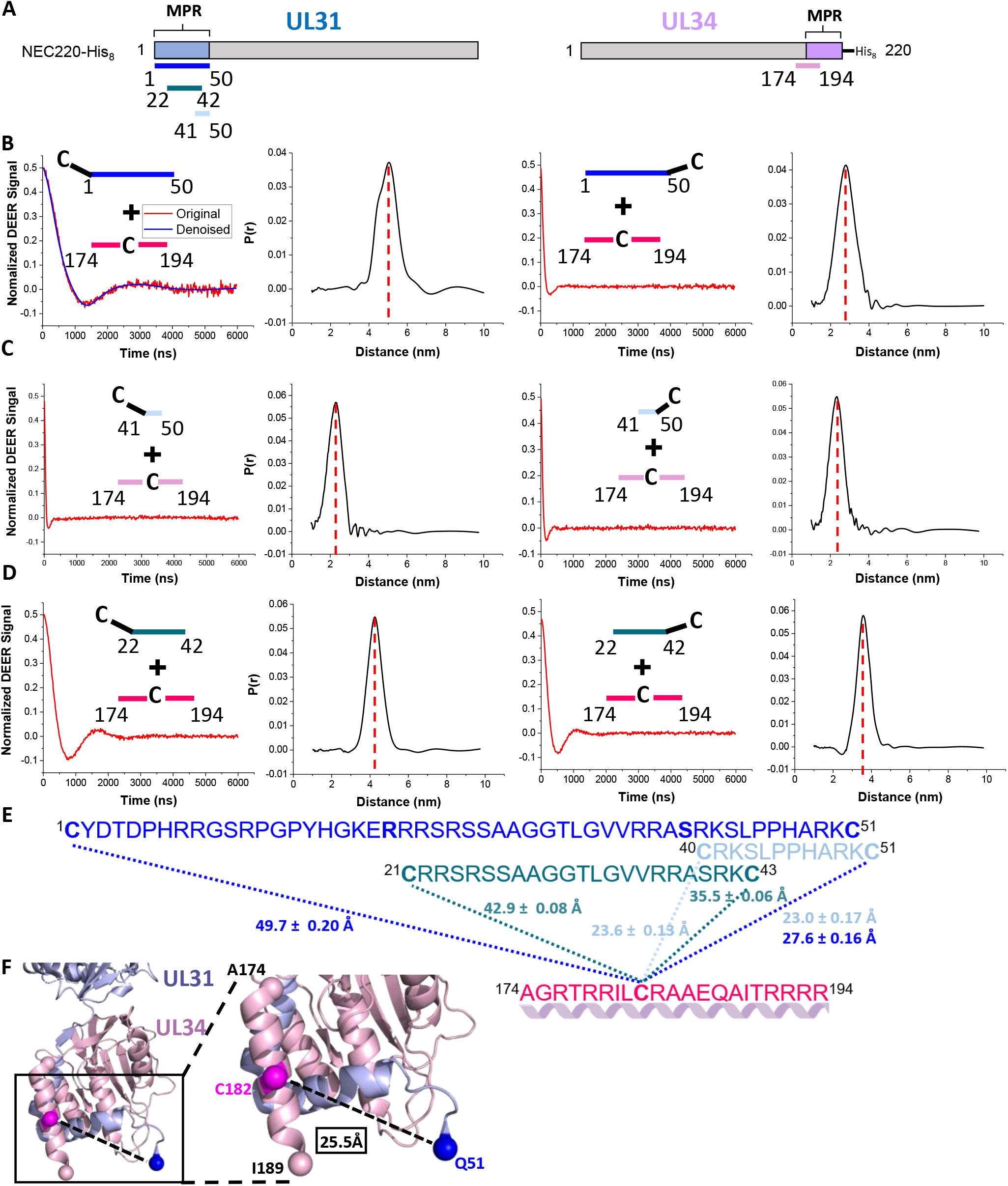
UL31 and UL34 MPR peptides interact in the presence of membranes. (A) Schematic depicting peptide location within the NEC sequence. (B, C, D and E) Representative experimental DEER data (left) and reconstructed interpeptide distance distributions (right) of at least two individual experiments. The “C” indicates the location of spin-labeled Cys. (B) UL31^(C1-50)^/UL34^(174-194)^ and UL31^(1-C51)^/UL34^(174-194)^, the blue line in the left panel is the denoised curve. (C) UL31^(C40-50)^/UL34^(174-194)^, UL31^(41-C51)^/UL34^(174-194)^, (D) UL31^(C21-42)^/UL34^(174-194)^, or UL31^(22-C43)^/UL34^(174-194)^. The peptides were mixed with POPC/POPS/POPA 3/1/1 SUVs in 1/200 P/L ratio. Each combination was repeated two times. (E) Combined sequences for each of the peptides used in DEER experiments with distances and associated standard error of the mean (68% confidence interval of the mean) for each tested probe location. UL31^(C1-C51)^ (royal blue), UL31^(C40-C51)^ (light blue), UL31^(C21-C43)^ (teal), UL34^(174-194)^ (pink). Bold letters denote probe location in different UL31 peptides. Predicted secondary structure for UL34 MPR shown in purple. (F) Bottom of homology modeled HSV-1 NEC onto PRV NEC. Inset shows distance measurement between Q51_UL31_ and C182_UL34_ shown as spheres. A174_UL34_ and I189_UL34_ shown as spheres. Images taken in PyMoL.

A common way to evaluate the DEER distance measurements is to compare them to the corresponding measurements in the high-resolution protein structures. Although the residues labeled in the DEER experiments were absent from the HSV-1 NEC structure (Bigalke &Heldwein, 2015), residues corresponding to 51-54 of UL31 and 177-189 of HSV-1 NEC were resolved in the crystal structure of the PRV NEC (Bigalke &Heldwein, 2015) and were modelled onto the HSV-1 NEC structure (Fig 7F). The distance between Q51_34_ (Cα) and C182_34_ (Cα) in the model was 25.5 Å (Fig 7F), which is similar to the DEER distances of 23.0 ± 0.17 Å and 27.6 ± 0.16 Å. The DEER results obtained with the MPR peptides are thus relevant to the MPRs in the context of the NEC.

### Chemical crosslinking confirms that UL31 and UL34 MPRs interact on membranes

To confirm UL31/UL34 MPR interactions identified by DEER, we performed chemical crosslinking. The UL31 peptides have a primary amine at K42_31_ and no sulfhydryls whereas the UL34 peptide has a sulfhydryl at C182_34_ and no primary amines, so the heterobifunctional SM(PEG)_6_ crosslinker that reacts with primary amines and sulfhydryls was used. SM(PEG)_6_, which can crosslink primary amines and sulfhydryls within 32.5 Å, should be capable of bridging the ~30-Å distance between K42_31_ and C182_34_ as measured by DEER. The UL31^(41–50)^/UL34^(174–194)^ and UL31^(22–42)^/UL34^(174–194)^ combinations were only crosslinked in the presence of SUVs whereas individually, UL31^(41–50)^ or UL34^(174–194)^ did not get crosslinked and UL31^(22–42)^ showed only low levels of crosslinking in the presence or absence of SUVs (S10 Fig), in agreement with the DEER data showing individual peptides do not form homodimers either in solution or on SUVs. The crosslinking results further establish that the peptides derived from the MPRs of UL31 and UL34 interact on the membranes.

### UL34 MPR peptide forms an α-helix in the presence of membranes

Circular dichroism (CD) (Kelly *et al*, 2005) was used to assess the secondary structure content of the UL31 and UL34 MPR peptides. A characteristic CD spectrum of an α helix has two negative troughs at 222 nm and 208 nm and a positive peak at 192 nm whereas the CD spectrum of a random coil has low ellipticity above 210 nm and negative values near 195 nm (Greenfield, 2006; Kelly *et al.*, 2005). The UL34^(174–194)^ was expected to form a helix because equivalent residues form α helices in the structures of PRV and HCMV UL34 homologs (Bigalke & Heldwein, 2015; Lye *et al*, 2015; Walzer *et al*, 2015; Zeev-Ben-Mordehai *et al*, 2015). UL34^(174–194)^ peptide adopted a random-coil conformation in solution but became α-helical in the presence of SUVs (S11B Fig), which suggested that its sequence has a propensity to form α-helical structure. By contrast, all UL31 peptides, UL31^(41–50)^, UL31^(22–42)^, or UL31^(1–50)^, adopted a random-coil conformation both in solution and in the presence of SUVs (S11C,E,G Figs). The spectra of the UL31^(41–50)^/UL34^(174–194)^ and UL31^(22–42)^/UL34^(174–194)^ combinations had helical signatures (S11DF Fig), but these were less pronounced than that of UL34^(174–194)^ alone (S11B Fig) whereas the spectrum of UL31^(1–50)^/UL34^(174–194)^ combination had no obvious helical signature (S11H Fig). We hypothesize that the helical signature of UL31/UL34 peptide combinations is due to UL34 and is less pronounced than that of UL34^(174–194)^ due to the UL34 signal being “diluted” by the unstructured UL31 peptides. The CD data suggest that the UL31 MPR is unstructured even in the presence of UL34 MPR and membranes.

## Discussion

Generation of membrane curvature lies at the core of the membrane budding ability of the NEC, but how the NEC accomplishes this is unclear. Previous work has shown that the NEC oligomerizes into hexagonal scaffold-like coats on the inner surface of budded vesicles (Bigalke *et al.*, 2014; Hagen *et al.*, 2015; Newcomb *et al.*, 2017) and that oligomerization is essential for budding both *in vivo* and *in vitro* (Bigalke &Heldwein, 2015; Bigalke *et al.*, 2014; Roller *et al.*, 2010). Membrane scaffolding is a common mechanism for generating both positive and negative membrane curvature, e.g., by the BAR domain proteins (reviewed in (Simunovic *et al*, 2019)) and HIV Gag (Schur *et al*, 2016), respectively. Therefore, one may conclude that formation of negative membrane curvature by the NEC could be driven by scaffolding alone. However, here we show that highly basic MPRs of the NEC are also required for budding and can alter lipid order by inserting into the lipid headgroups. Therefore, we hypothesize that the NEC-mediated membrane budding is driven by a mechanism that combines scaffolding with lipid insertion. Furthermore, we show that the NEC can generate negative Gaussian curvature required for the formation and scission of the bud neck, which is consistent with the NEC-induced scission observed *in vitro* (Bigalke *et al.*, 2014). The NEC is thus a self-contained membrane-budding machine capable of completing multiple actions in the budding process, at least, *in vitro*.

### Electrostatic forces govern NEC/membrane interactions

Previously, we showed that the NEC MPRs are necessary for the membrane recruitment of the soluble NEC *in vitro* through electrostatic interactions (Bigalke *et al.*, 2014). Electrostatic interactions between basic residues and acidic lipids commonly serve to recruit cytoplasmic proteins to membranes (Mulgrew-Nesbitt *et al.*, 2006), but the NEC is anchored in the INM through the TM helix of UL34 (Schuster *et al.*, 2012), which left uncertain the role of the MPRs in membrane budding. Here, we found that the MPRs – especially, the UL31 MPR – are necessary for membrane budding and can induce lipid ordering. Both phenomena require basic clusters within the UL31 MPR. Basic clusters govern membrane interactions of proteins such as Src kinase (Sigal *et al*, 1994), myristoylated Alanine-Rich C-Kinase Substrate (MARCKS) (Kim *et al*, 1994a), neuromodulin (Kim *et al.*, 1994a), the BAR domain proteins (Itoh *et al*, 2005; Mulgrew-Nesbitt *et al.*, 2006; Peter *et al*, 2004), and HIV Gag (Zhou *et al*, 1994). Moreover, it has been proposed that interactions of basic clusters with the membrane could promote negative membrane curvature by concentrating negatively charged lipids within the membrane (Bassereau *et al*, 2018). We hypothesize that interactions between the basic clusters within the UL31 MPR and the membrane drive formation of negative membrane curvature by the HSV-1 NEC. Considering that basic clusters are found in the MPRs of many UL31 homologs (S1 Fig), their involvement in curvature formation may be a conserved feature of the NEC budding mechanism across different herpesviruses.

In addition to basic residues, the HSV-1 UL31 MPR contains six serines that are phosphorylated by the US3 viral kinase (Chang & Roizman, 1993; Kato *et al.*, 2005; Mou *et al.*, 2009). The role of UL31 phosphorylation in nuclear egress is unclear, but phosphomimicking serine-to-glutamate mutations of these six serines inhibits nuclear egress and HSV-1 replication (Mou *et al.*, 2009), suggesting that phosphorylation may inhibit nuclear egress, by an unknown mechanism. Here, we observed that serine-to-glutamate mutations within the UL31 MPR blocked NEC-mediated budding *in vitro*. Glutamates, just as phosphates, are negatively charged, and since NEC/membrane interactions require a sufficiently high net positive charge of the UL31 MPR, introducing negative charges would disrupt proper NEC/membrane interactions. Indeed, phosphorylation and phosphomimicking mutations decrease protein/membrane interactions of the F-BAR domain of syndapin I (Quan *et al*, 2012), MARCKS (Kim *et al.*, 1994a; Kim *et al*, 1994b), neuromodulin (Kim *et al.*, 1994a), Cdc15 (Roberts-Galbraith *et al*, 2010), PTEN (Das *et al*, 2003), and dynamin I (Powell *et al*, 2000). Therefore, we hypothesize that phosphomimicking mutations block capsid nuclear egress by reducing the net positive charge of the UL31 MPR thereby inhibiting NEC-mediated budding. Phosphorylation also introduces negative charges and would have a similar inhibitory effect on budding. We speculate that HSV-1 uses phosphorylation to inhibit the membrane-budding activity of the NEC and, thus, nuclear egress, by fine-tuning its membrane interactions. In this way, phosphorylation could serve as an “off” switch that prevents unproductive membrane budding prior to the arrival of the capsid.

### Lipid ordering by MPR insertion in combination with scaffolding generates negative mean curvature for the growing bud

NEC-mediated membrane budding proceeds through two distinct steps: formation of the bud and scission of the bud neck. Bud formation requires generation of the negative mean membrane curvature. The two most common mechanisms of curvature generation, be it positive or negative, are peripheral insertion of protein into lipid bilayers and scaffolding of the curvature by protein oligomers (reviewed in (Campelo *et al*, 2010; Campelo *et al*, 2008b; McMahon &Boucrot, 2015; Zimmerberg & Kozlov, 2006). We propose that NEC-mediated membrane budding is driven by a mechanism that combines scaffolding with insertion. Previous studies already established that the NEC oligomerizes into hexagonal scaffold-like coats on the inner surface of budded vesicles (Bigalke *et al.*, 2014; Hagen *et al.*, 2015; Newcomb *et al.*, 2017), and this oligomerization is essential for budding both *in vivo* and *in vitro* (Bigalke & Heldwein, 2015; Bigalke *et al.*, 2014; Roller *et al.*, 2010). Therefore, formation of negative membrane curvature by the NEC requires membrane scaffolding by NEC oligomers. Here, we demonstrated that highly basic MPRs of the NEC are also required for budding and can induce ordering of lipid headgroups and upper acyl chain regions in the protein-proximal leaflet of the membrane bilayer.

Lipid ordering is mediated by the UL31 MPR that engages membranes directly by inserting into the lipid headgroups. But while many peripheral membrane proteins use amphipathic helices for membrane interactions, the UL31 MPR maintains a random-coil conformation even in the presence of membranes. Therefore, we think that the basic clusters within the UL31 MPR form fingertip-like projections that interact with the lipid headgroups in a multidentate manner (Fig 8), similarly to the membrane-interacting fusion loops (FLs) of class II viral fusogens, in which three or six FLs ensure sufficient grip on the membrane (Modis, 2014). It is unclear how many residues in the UL31 MPR insert into the membrane; however, given the low volume of protein detected in the membrane by NR, relatively few residues are involved. Ordering of the upper acyl chains could be due to the insertion of the UL31 MPR into the upper acyl chain region. Alternatively, the UL31 MPR could be drawing the headgroups together and constraining the motion of the upper acyl chains and thus the spin label located there, without directly occupying the upper acyl chain region.

**Fig. 8.**
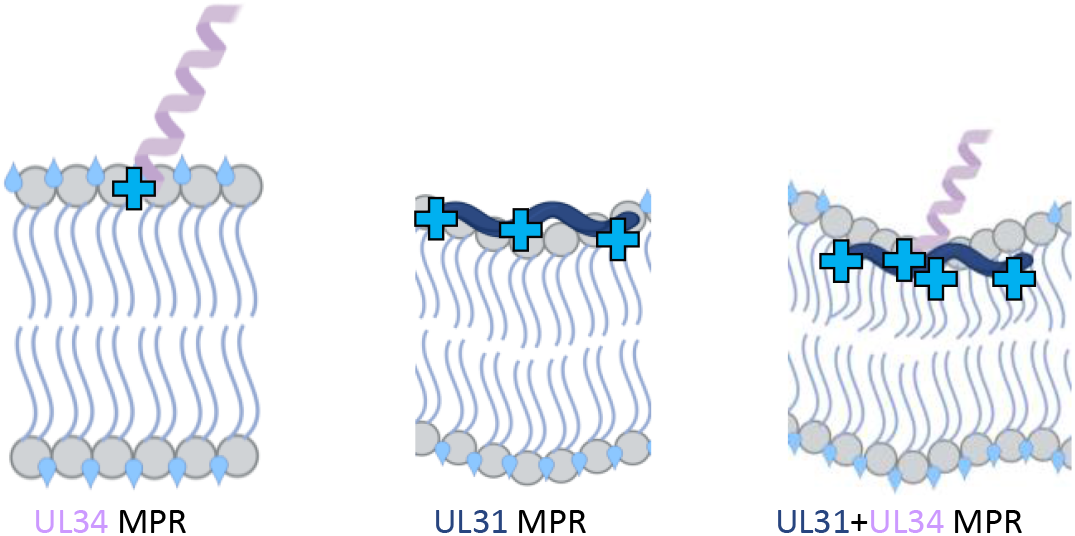
Model of negative mean membrane curvature and negative Gaussian curvature generation by NEC MPR peptide-membrane interactions. The UL34 MPR peptide (light purple) with a C-terminal patch of basic residues (light blue cross) alone is insufficient to drive ordering of lipid headgroups and acyl chain region nor displace water (light blue tear drops). The UL31 MPR peptide (dark blue) alone can induce ordering of the lipid headgroups, accompanied by outer leaflet dehydration. Combination of the UL31 and UL34 MPR peptides results in both lipid headgroup and upper acyl chain ordering along with membrane dehydration, resulting in the generation of local negative mean curvature in the protein-proximal leaflet. All images created with BioRender.com.

The cooperative effect of the UL31 and UL34 MPR peptides on lipid ordering as well as a greater depth of insertion of UL31 MPR peptides in the presence of UL34 MPR peptide suggest that the UL31 and the UL34 MPR peptides interact in the presence of the membrane, which we detected by both DEER and chemical crosslinking. Whereas the UL31 MPR interacts with the membrane directly, the UL34 MPR likely assists in positioning the UL31 MPR for optimal penetration necessary to induce the required degree of lipid ordering (Fig 9). The HSV-1 UL34 MPR is predicted to form an α helix and, indeed, becomes α-helical in the presence of the membrane. Although this region was unresolved in the HSV-1 NEC crystal structure (Bigalke &Heldwein, 2015), the corresponding region in HCMV (Lye *et al.*, 2015; Walzer *et al.*, 2015) and PRV (Bigalke &Heldwein, 2015; Zeev-Ben-Mordehai *et al.*, 2015) homologs forms an α helix oriented perpendicularly to the membrane. To reflect this, were have modeled the UL34 MPR peptide such that its α-helical segment is oriented perpendicularly to the membrane, which positions its basic cluster to interact with the membrane and, presumably, with the UL31 MPR (Fig 8).

**Fig. 9.**
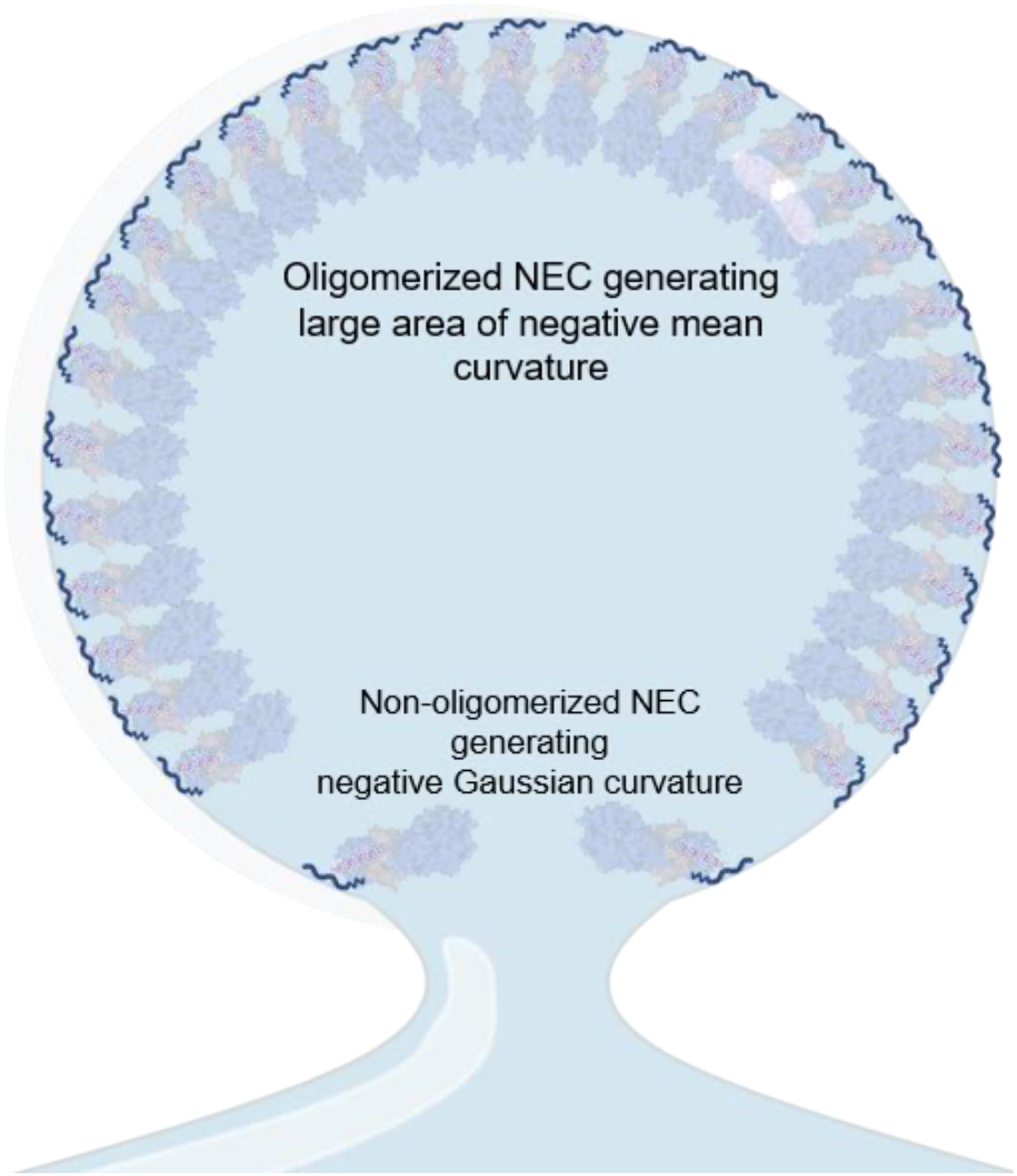
Model of negative mean membrane curvature and negative Gaussian curvature generation by NEC MPR peptide-membrane interactions. Oligomerized NEC forces MPRs to work in concert and generate larger areas of negative mean curvature (top inset). Non-oligomerized NEC adopts a more flexible orientation and the MPRs generate negative Gaussian curvature to perform scission (bottom inset). All images created with BioRender.com.

It has been proposed that protein-mediated ordering of the lipid headgroups results in dehydration of the protein-proximal leaflet leading to tighter lipid packing and shrinking of the local area (Ge &Freed, 2009), leading to the formation of negative mean membrane curvature. On a flat substrate, this would result in membrane thickening, and, indeed, the NR experiments revealed a thickening of the tethered bilayer after incubation with NEC220. Therefore, we hypothesize that the MPR-induced ordering of lipid headgroups and upper acyl chains generates negative mean membrane curvature. Given that MPR/membrane interactions would generate curvature only locally, we hypothesize that generation of negative mean membrane curvature over a large membrane area requires NEC oligomerization into a hexagonal scaffold. As the membrane-tethered NEC heterodimers oligomerize into the hexagonal scaffold, they can create compressive pressures that would generate negative mean curvature, driving vesicle budding (Kim &Sung, 2001; Stachowiak *et al*, 2013). In this manner, the lipid ordering and oligomerization work together to mold the associated membrane into a spherical shape.

### NEC can achieve scission by generating negative Gaussian curvature

In addition to generating membrane buds, the NEC can also drastically constrict the necks of the budded vesicles via membrane remodeling *in vitro* (Bigalke *et al.*, 2014). We found that UL31 and UL34 MPR peptides can generate NGC, which is the type of curvature topologically required for formation of the scission neck, and that their effect on NGC formation is cooperative, which parallels their effect on lipid ordering observed by the ESR. Based on the quantitative measurements of NGC in the MPR-induced lipid phases, we estimate that the NEC could generate the membrane scission neck with an inner diameter of 7.2 nm, which is consistent with the diameters of the necks formed by mitochondrial fission proteins capable of spontaneous scission (Lee *et al.*, 2017) and with the theoretical calculations (Kozlovsky &Kozlov, 2003). The ability of the MPRs to generate tight membrane curvatures found in scission necks in other biological systems suggest that they contribute likewise to NEC-induced bud scission observed *in vitro* (Bigalke *et al.*, 2014; Lorenz *et al.*, 2015b).

While NEC demonstrates an intrinsic membrane scission ability *in vitro* (Bigalke *et al.*, 2014; Lorenz *et al.*, 2015b), efficient nuclear egress at least in some cell types (Arii *et al.*, 2018) if not in others (Crump *et al.*, 2007) requires ESCRT-III machinery. Several enveloped viruses, notably HIV, recruit cellular endosomal sorting complexes required for transport III (ESCRT-III) (reviewed in (Alonso *et al*, 2016; Hurley, 2015; Hurley & Cada, 2018; McCullough *et al*, 2018; Votteler & Sundquist, 2013)) to mediate scission during viral budding. ESCRT-III proteins accomplish scission of the bud neck by forming a spiral polymer on the inward face of the neck and constricting it (Effantin *et al*, 2013; Nguyen *et al*, 2020). Not all enveloped viruses, however, recruit ESCRT-III proteins for membrane scission. For example, Influenza A virus deploys the amphipathic helix within its M2 channel (Rossman *et al*, 2010) (reviewed in (Rossman & Lamb, 2011, 2013)), which has been proposed to mediate neck scission through a mechanism that involves the generation of the NGC (Schmidt *et al*, 2013a).

The neck generated by the NEC may not be sufficiently narrow to trigger spontaneous membrane scission with high enough efficiency required for vesicle release in the context of HSV-1 nuclear budding. If so, low efficiency of this NEC-mediated scission could, in principle, account for the need to recruit ESCRT-III machinery to increase the efficiency of membrane bud scission during nuclear egress, in a cell-type-specific manner. This scenario is reminiscent of Ebola virus, where the viral matrix protein VP40 mediates membrane budding *in vitro* (Soni &Stahelin, 2014) yet recruits ESCRT-III machinery *in vivo* (Licata *et al*, 2003; Silvestri *et al*, 2007) (reviewed in (Gordon *et al*, 2019)). Future experiments will address the coordination of efforts between the NEC and the ESCRT-III proteins in mediating nuclear egress.

### A model of membrane curvature generation by the NEC

While the NEC can form both negative mean curvature and NGC, it is yet unknown what determines the transition from dome formation to neck formation and scission. We postulate that this switch depends on the oligomeric state of the NEC. Within the hexagonal lattice, the NEC heterodimers adopt vertical orientations (Bigalke *et al.*, 2014; Hagen *et al.*, 2015), yet the NR measurements suggest that a significant fraction of the NEC may have tilted or flat orientations. Therefore, we hypothesize that on the membrane, there are regions with high and low densities of NECs. At higher NEC densities, oligomerization would promote an upright orientation whereas at lower NEC densities, individual NEC heterodimers would experience greater orientational freedom.

Putting together our experimental observations, we propose the following model of curvature generation by the NEC (Fig 9). We hypothesize that in areas with high NEC density, such as the body of the budding vesicle, the NEC oligomerizes into the hexagonal scaffold. While the MPRs of the NECs have the capacity to generate NGC, in the body of the bud, the hexagonal scaffold forms a rigid frame that constrains the membrane into a defined spherical architecture, promoting negative mean curvature. As more NECs are recruited and oligomerize, the hexagonal scaffold expands, and the budding vesicle grows. However, at membrane regions not covered by the hexagonal coat, such as near the rim of the bud, NECs may mainly exist as unconstrained individual heterodimers or, perhaps, individual hexamers. At these regions, membrane interactions by individual NEC heterodimers could facilitate the induction and stabilization of NGC to produce saddle-shaped deformations necessary for scission (Fig 9). Future experiments will address how membrane interactions by the MPRs in the context of the full NEC are coordinated with NEC oligomerization to bring about membrane budding.

## Methods

### Cloning of expression constructs

Cloning of constructs encoding HSV-1 strain F UL31 with boundaries 1-306, 41-306 and 51-306 is described elsewhere (Bigalke *et al.*, 2014). Primers used for cloning are listed in S3 Table. Site-directed mutagenesis of the UL34 mutant with boundaries 1-185 was performed by restriction digest and ligation with SalI and NotI. Site-directed mutagenesis of the UL31 mutant with boundaries 1-306 (S11E/S24E/S43E) was performed by two rounds of inverse PCR on a full-length UL31 that already contained an S43E mutation and 24 blunt-end ligation. S11E/S24E/S26E/S27E/S40E/S43E was generated by three sequential inverse PCR and blunt-end ligation reactions. The first round was to generate S11E followed by blunt-end ligation and inverse PCR to generate S11E/S24E/S26E/S27E, followed by blunt-end ligation and inverse PCR to generate S11E/S24E/S26E/S27E/S40E/S43E. Site-directed mutagenesis of UL31 mutants with boundaries 41-306 (R49S/K50S, R41S/K42S/R49S/K50S, L44A, S43E, R41S/K42S/S43R/L44K/R49S/K50S, R41S/K42S/P45R/P46K/R49S/K50S, R41S/K42S/H47R/A48K/R49S/K50S, P45A/P46A, ^41^KSPKLHRARP^50^) was performed by inverse PCR. Site-directed mutagenesis for the UL31 mutant with boundaries 41-306 and mutations R41S/K42S was performed by restriction digest and ligation with BamHI and NotI. Site-directed mutagenesis of UL31 mutant with boundaries 47-306 containing mutations H47R/A48K was performed by inverse PCR.

A gene block with the codon-optimized DNA sequence for UL31 residues 1-50 was purchased from Integrated DNA Technologies (IDT). PCR was performed on the gene block with primers containing the restriction digest sites for BamHI and NotI. The resulting PCR product was purified and digested with BamHI and NotI and ligated into pGEX-6P-1 with an N-terminal GST tag for solubility and affinity purification purposes. Inverse PCR followed by blunt end ligation was used to develop UL31^(C1-50)^ and UL31^(1-C51)^. Three rounds of inverse PCR and blunt end ligation were needed for UL31^(C1-50)^ due to introduction of point mutations.

All constructs generated in this work are listed in S3 Table.

### Protein purification

Plasmids encoding HSV-1 UL31 and UL34 were co-transformed into *E. coli* LOBSTR-BL21 (DE3) cells and expressed at 25 °C for 16 hours after lactose-derived autoinduction (Studier, 2005). Cells were resuspended in lysis buffer (50 mM HEPES pH 7.5, 500 mM NaCl, 0.5 mM TCEP, 10% glycerol) in the presence of Complete protease inhibitor (Sigma-Aldrich) and lysed using a M-110S microfluidizer (Microfluidics). Cell lysate was spun down at 12,500 rpm in a Beckman J2-21 floor centrifuge. All purification steps were performed at 4 °C. The clarified cell lysate was first passed over Ni-NTA Sepharose resin (GE Healthcare). The resin was washed with wash buffer (lysis buffer containing 20-40 mM imidazole). Bound protein was eluted with elution buffer (lysis buffer containing 250 mM imidazole) and loaded onto glutathione Sepharose resin (GE Healthcare) to separate bound NEC from excess His_6_-SUMO-UL31. After washing with lysis buffer containing 1 mM EDTA, His_6_-SUMO and GST tags were cleaved on the glutathione Sepharose column for 16 hours using PreScission protease produced in-house using a GST-PreScission fusion protein expression plasmid. Cleaved NEC and His_6_-SUMO were eluted from the glutathione sepharose column with lysis buffer and diluted to 100 mM NaCl with 50 mM HEPES pH 7.5, 0.5 mM TCEP, 10% glycerol. NEC was separated from His_6_-SUMO using a cation exchange resin (HiTrap SP XL, GE Healthcare) with a 200 mM to 600 mM NaCl gradient in 20 mM HEPES pH 7.0, 0.5 mM TCEP. Each NEC construct was purified to homogeneity as assessed by 12% SDS-PAGE and Coomassie staining. Fractions containing NEC were diluted to 100 mM NaCl with 20 mM HEPES pH 7.0, 0.5 mM TCEP and concentrated up to ~1.5 mg/mL and stored at −80 °C to avoid aggregation and degradation at 4°C. Protein concentration was determined by absorbance measurements at 280 nm. The typical yield was 0.5 mg per L TB culture. UL31 1-50 peptides UL31^(1-50)^, UL31^(C1-50)^, and UL31^(1-C51)^ were expressed and purified with the same buffers. Briefly, these three UL31 MPR peptides were passed over glutathione Sepharose resin and washed with lysis buffer containing 1 mM EDTA. The GST tag was cleaved as outlined above. The glutathione Sepharose column eluate was then concentrated to 500 μL and passed over an S75 10/300 size exclusion column (GE healthcare) with 20 mM HEPES pH 7.0, 100 mM NaCl, 0.5 mM TCEP. Fractions containing UL31 MPR peptides were concentrated up to 5 mg/mL and stored at −80 °C.

### Liposome preparation

Liposomes were prepared as described previously (Bigalke *et al.*, 2014). Briefly, MLVs were made by mixing 1-palmitoyl-2-oleoyl-sn-glycero-3-phosphocholine (POPC), 1-palmitoyl-2-oleoyl-sn-glycero-3-phospho-L-serine (POPS) and 1-palmitoyl-2-oleoyl-sn-glycero-3-phosphate (POPA) (Avanti Polar Lipids) in a molar ratio of 3/1/1 POPC/POPS/POPA, followed by vacuum drying the mixture and resuspending in 200 μL 20 mM HEPES pH 7.0, 100 mM NaCl, 0.5 mM TCEP to shake for 0.5 hours in a 37 °C incubator (Bigalke *et al.*, 2014; Zhao *et al*, 2010). The lipid mixture was then vortexed and used immediately. For GUVs, lipids were mixed in a molar ratio of 58% POPC/11% POPE/9% POPA/9% POPS/5% cholesterol/5% DGS-NTA/3% POPE Atto594, of which 5 μL was spread on the surface of an ITO-covered slide and vacuum desiccated for 30 minutes. A vacuum-greased O-ring was placed around the dried lipid mixture and the VesiclePrep Pro (Nanion Technologies) was used to produce an AC field (sinusoidal wave function with a frequency of 8Hz and amplitude 2V) before adding 270 μL of lipid swelling buffer (300mM sucrose dissolved in 5 mM Na-HEPES, pH 7.5). A second ITO-covered slide was used to cover the lipid/buffer mixture after 3 minutes followed by a 2-hour swell and a 5-minute fall step. GUVs were used immediately and diluted 1/20 with 20 mM HEPES pH 7.0, 100 mM NaCl, 0.5 mM TCEP.

### Membrane co-sedimentation assay

3 μg of protein was incubated with or without 15 μg freshly prepared MLVs (as detailed above) at 20 °C for 30 minutes. The samples were centrifuged at 16,000 g for 20 minutes at 4 °C. Aliquots of protein/MLV pellet and protein supernatant were analyzed by 12% SDS-PAGE and Coomassie staining. The amount of protein that pelleted with MLVs was determined by densitometry analysis of gels imaged using a LiCor Odyssey CLx imager and quantified using imageJ. For each protein, band intensities of the pelleted protein were integrated and expressed as a percentage of the total integrated intensity of protein bands in the pelleted sample and supernatant sample. Background levels of pelleted protein in the absence of vesicles were subtracted from levels of protein sedimentation in the presence of vesicles. Each experiment was done with duplicate technical replicates with at least three biological replicates and the average value and standard error of the mean is reported. Data were plotted using GraphPad Prism 9.0.

### GUV budding assay

Fluorescently labeled giant unilamellar vesicles (GUVs) were co-incubated for 3 minutes with the soluble NEC and the membrane impermeable dye, Cascade Blue Hydrazide (ThermoFisher Scientific). The GUV contained 18% negatively charged lipids (58% POPC/11% POPE/9% POPA/9% POPS/5% cholesterol/5% DGS-NTA/3% POPE Atto594), which closely resembles the inner nuclear membrane (Keenna *et al*, 1970; Neitcheva &Peeva, 1995). Budding events manifested as the appearance of intraluminal vesicles (ILVs) containing Cascade Blue within the GUVs (Fig 1C). 5 μL of the above GUV composition and 2 μL Cascade Blue Hydrazide were mixed with a final concentration of 1.5 μM NEC for a total volume of 100 μL. Each sample was visualized using a Nikon A1R Confocal microscope. Background levels of intraluminal vesicles were counted in the absence of NEC and subtracted from counts of intraluminal vesicles in the presence of NEC. Experiments were performed with at least 3 technical replicates and at least 3 biological replicates. All counts were normalized to NEC220-His_8_ budding. Levels of budding are broken down into three categories based on statistical significance, poor (0-49%, p<0.0005=***), moderate (50-74%, p<0.05=*, p<0.005=**) and efficient (75-100%). The NEC220∆50-RKRK-His_8_ and NEC220∆40-P45A/P46A-His_8_ mutants were exceptions because they supported efficient budding (85% and 75%, respectively), yet the p value was <0.05. The standard error of the mean is reported from at least three individual experiments. Data was plotted using GraphPad Prism 9.0.

### Coflotation assay

NEC sensitivity to membrane curvature was tested using coflotation as described previously (Silverman *et al*, 2012). Briefly, 1.5 μg NEC was incubated with or without large unilamellar vesicles (LUVs) [POPC, POPS, and POPA mixed in a 3/1/1 molar ratio as previously described (Bigalke *et al.*, 2014)] at room temperature for 20 minutes in 50 mL PBS. KCl was added to 200 mM concentration to reduce nonspecific protein–membrane interactions, and samples were incubated for 15 minutes at room temperature. Optiprep (Sigma) was added to a final concentration of 30% in a 500 mL volume. Samples were placed at the bottom of a 5-mL centrifugation tube (Beckmann) and overlaid with 4 mL 15% Optiprep and 500 mL 3% Optiprep in PBS. The samples were next centrifuged in a Beckman SW-55 Ti rotor at 246,000 g for 3 hours at 4 °C, and 1 mL fractions were collected beginning at the top. Protein was precipitated with 20% trichloroacetic acid for 30 minutes on ice. Sample was washed with 750 μL cold acetone and then spun in a tabletop centrifuge for 10 minutes at 14,000 rpm. This was repeated for a total of 3 washes. Samples were then analyzed by western blot for UL31 as previously described (Bigalke *et al.*, 2014). The standard error of the mean is reported from at least two individual experiments. Data was plotted using GraphPad Prism 9.0.

### Neutron Reflectometry

Silicon wafers (100, n-doped to a conductivity of 1-100 Ω cm) of 5 mm thickness and 75 mm diameter were coated with 40 Å Cr, for adhesion purposes, followed by 140 Å Au by magnetron sputtering on a Denton Vacuum Discovery 550 Sputtering System at the NIST Center for Nanoscale Science and Technology cleanroom. The substrate was then immersed for 8 hours in an ethanolic solution of the thiol-lipid linking molecule HC18 (Z20-(Z-octadec-9-enyloxy)-3,6,9,12,15,18,22-heptaoxatetracont-31-ene-1-thiol)) (Rakovska *et al*, 2015) and βME (β-mercaptoethanol) in a 3/7 molar ratio and a total concentration of 0.2 mM (M = mol/L). The resulting self-assembled monolayer (SAM) was rinsed in ethanol and dried in a nitrogen stream. The coated surface of the sample wafer was mounted facing a 100 μm reservoir defined by a 65-mm inner diameter cylindrical Viton gasket separating the sample wafer from a rough backing wafer (Eells *et al*, 2019). The backing wafer was perforated by single inlets and outlets, which were coupled by IDEX Health and Science (Oak Harbor, WA) flat-bottomed fittings to external tubing for solution exchanges, which were performed using at least 7.5 mL flowing at 2.5 mL/min. To prepare multilamellar vesicles (MLVs), a solution of POPC/POPS/POPA in a 3/1/1 molar ratio was prepared at 10 mg/mL in 1 M NaCl, subjected to 40 minutes of bath sonication, and injected into the sample cell. Incubation proceeded for at least 1.5 hours, followed by flushing with pure water to lyse the vesicles via osmotic stress, forming a sparsely tethered lipid bilayer membrane.

NR experiments were carried out on the MAGIK vertical reflectometer (Dura *et al*, 2006) at the National Institute for Standards and Technology (NIST) Center for Neutron Research (NCNR). A monochromatic beam of wavelength λ=5.000 Å impinged on the interface between the coated surface of the sample wafer and the liquid in the sample cell reservoir. The pre-sample collimating slits were chosen to maintain a constant illuminated interface area for each measured angle θ. The post-sample collimation was chosen to allow the entire reflected beam to impinge on the detector, which was positioned at an angle 2θ relative to the incoming beam direction to measure specular reflection. Each reflectivity curve covered a range in scattering wavevector Q=4πλ^-1^sin(θ) from 0.008 Å^-1^ to 0.251 Å^-1^.

The reflectivity was calculated as R(Q)=(I(Q)-I_B_(Q))/I_0_(Q). Here I(Q) is the measured count rate (normalized to a much larger monitor count rate to account for fluctuations in beam intensity) at the specular condition. I_B_(Q) is the background intensity, which arises primarily from incoherent scattering from the liquid reservoir and is calculated by linear interpretation of the intensities measured with the detector at off-specular positions 1.5θ and 2.5θ. I_0_(Q) is the incident beam intensity and is directly measured through the silicon substrate at θ=0 with the detector positioned in line with the incident beam.

NR data were analyzed using the composition space modeling procedures described previously (Shekhar *et al*, 2011). Briefly, the composition space model arranges the known molecular components of the tethered bilayer and protein at the substrate surface; any unfilled space is assumed to be filled with water. Because the neutron scattering length density (nSLD) of each component is known or can be estimated from its elemental composition and molecular volume, an average nSLD profile can be calculated as a function of distance from the substrate surface. This nSLD profile in turn corresponds to a predicted R(Q) which can be optimized to the experimental data, using as parameters the spatial arrangement of the molecular components. Replacing all H_2_O in the membrane-bathing buffer with D_2_O provides contrast and allows unambiguous determination of the nSLD profile associated with both measured R(Q) curves by simultaneous optimization of the two contrast conditions (Kirby *et al*, 2012).

The protein profile was parameterized in two ways for comparison. The Catmull-Rom spline, or “freeform” profile, makes no assumptions about the shape of the volume occupied by the protein, but does assume that the nSLD of the protein is equal to its average value for the entire protein. Alternatively, an “orientation” profile is used, in which the protein profile is calculated from the crystallographic structure of the NEC complex (PDB: 4ZXS) (Bigalke & Heldwein, 2015) rotated by Euler angles α and β, with the volume of the MPRs represented in the appropriate molar ratio by a smoothed box function. The Euler angles are defined in a *x-y-z* extrinsic rotation scheme, where the *z* axis is co-directional with the surface normal. Because NR is sensitive to the nSLD only in the *z* direction, it is not sensitive to the final rotation about the *z* axis, γ. Each profile was convolved with a width 4.1 Å Gaussian function to account for surface roughness. The orientation models require fewer parameters than freeform models and do account for spatial variations in nSLD, but assumes a single, rigid structure for the protein.

Optimization was performed on the Bridges (Nystrom *et al*, 2015; Towns *et al*, 2014) high performance computing system using the DREAM Markov Chain Monte Carlo (MCMC) algorithm (Vrugt *et al*, 2009) implemented in the software package Refl1D (Kienzle *et al*, 2016). Confidence intervals (CI) on parameters and model predictions were calculated from parameter distributions derived from 14.4 million DREAM samples after the optimizer had reached steady state.

### Lipid preparation for electron spin resonance

POPC/POPS/POPA were mixed at a 3/1/1 molar ratio with 0.5% (mol/mol) of spin-labeled lipid in chloroform and dried under N_2_ gas. The dried mixture was placed under vacuum overnight to remove any remaining chloroform. To prepare SUVs, dried lipids were resuspended in pH 7.0 buffer (20 mM HEPES, 100 mM NaCl, 0.5 mM TCEP) and sonicated in an ice bath for at least 20 minutes or until the solution became clear. The SUV solution was then subject to ultracentrifugation at 13,000 rpm for 10 minutes for further clarification to remove the possible membrane debris.

### UL31 and UL34 MPR peptides

Peptides UL31^(41-50)^, UL31^(C40-50)^, UL31^(41-C51)^, UL31scr^(41-C51)^, UL31^(C40-50 R41S/K42S)^, UL31^(22-42)^, UL31^(C21-42)^, UL31^(22-C43)^, UL31scr^(22-C43)^, UL34^(174-194)^, and UL34^scr(174-194)^ were purchased from Peptide 2.0. All peptides were N-terminally acetylated and C-terminally amidated and ≥ 95% pure.

### Peptide labeling

For peptide labeling, desired amounts of UL31 or UL34 peptides were dissolved in pH 8.0 buffer (5 mM HEPES, 10 mM MES, 150 mM NaCl) and mixed with 10-fold excess MTSL (S-(2,2,5,5-tetramethyl-2,5-dihydro-1H-pyrrol-3-yl) methyl methanesulfonothioate) dissolved in ethanol (200 mM); the volume of the ethanol added was less than 5% of the total volume. The mixtures were kept overnight in the dark at RT as previously described (Lai & Freed, 2014). The spin-labeled peptides were then subjected to purification using FPLC with a GE Superdex Peptide 10/300 GL at a flow rate of 0.2 mL/min for 150 minutes. The fractions containing the peptides were lyophilized overnight and dissolved in pH 7.0 buffer (20 mM HEPES, 100 mM NaCl, 0.5 mM TCEP).

### Continuous wave ESR (CW-ESR) on lipid probes

The desired amount of peptide and SUVs (3/1/1 POPC/POPS/POPA molar ratio) were mixed at RT for 30 minutes. The final amount of the lipid in each sample was 1 mg. The ESR spectra were collected on an ELEXSYS ESR spectrometer (Bruker Instruments, Billerica, MA) at X-band (9.5 GHz) at 25 °C using an N_2_ Temperature Controller (Bruker Instruments, Billerica, MA). The ESR spectra from the labeled lipids were first denoised whenever necessary (Srivastava *et al*, 2016). They were then analyzed using the NLLS fitting program based on the stochastic Liouville equation (Budil *et al*, 1996; Liang & Freed, 1999) using the Microscopic Order Macroscopic Disorder (MOMD) model as in previous studies (Ge & Freed, 2003, 2009, 2011; Ge *et al*, 2001; Smith & Freed, 2009). The A and g values of the spins are determined using the low temperature ESR measurements. Two sets of parameters that characterize the rotational diffusion of the nitroxide radical moiety in spin labels are generated. The first set consists of R_⊥_ and R_∥_, which are respectively the rates of rotation of the nitroxide moiety around a molecular axis perpendicular and parallel to the preferential orienting axis of the acyl chain. The second set consists of the ordering tensor parameters, S_0_ and S_2_, which are defined as follows: S_0_=<D_2,00_>=<1/2(3cos^2^θ-1)>, and S_2_=<D_2,02_+D_2,0-2_> =<√(3/2)sin^2^θcos2φ>, where D_2,00_, D_2,02_, and D_2,0-2_ are the Wigner rotation matrix elements and θ and φ are the polar and azimuthal angles for the orientation of the rotating axes of the nitroxide bonded to the lipid relative to the director of the bilayer, i.e., the preferential orientation of lipid molecules (Ge & Freed, 2009; Liang & Freed, 1999), with the angular brackets implying ensemble averaging.

S_0_ indicates the strength of the alignment of the chain segment to which the nitroxide is attached along the normal to the lipid bilayer, which is correlated with hydration/dehydration of the lipid bilayers (Ge & Freed, 2003). S_2_ is the measurement of the molecular non-axiality of the motion of the spin label. It was found to be much smaller than S_0_, with much less sensitivity to changes in bilayer structure in our studies. Therefore, S_0_ is the more important parameter for this study. The estimated error of S_0_ from the NLLS fit for the spectra (the typical standard deviation obtained in the fitting) is about ±0.005-0.008 from at least three individual experiments.

### Vesicle sedimentation assay for partition ratio

Sucrose-loaded LUVs (3/1/1 POPC/POPS/POPA molar ratio) were prepared as described previously (Buser & McLaughlin, 1998; Kuo *et al*, 2011). Approximately 100 μM peptide was incubated with 10 mM LUVs in a 1/1 ratio for 1 hour at 37 °C. The final lipid concentration was confirmed by a phosphate assay (Ames, 1966). The mixtures were then centrifuged at 100,000 × g for 1 hour at 25 °C. The pellets were washed briefly before ESR measurement. The amount of the spin-labeled peptides in the supernatant and the pellets were determined by CW-ESR using the build in double integration tool in the Bruker XEPR program. The partition ratio for peptide is defined as the amount of peptide in the pellet to the amount of peptide in the supernatant. Data shown is from three independent experiments and the standard deviation is reported.

### Power saturation CW-ESR

The spin-labeled peptides were mixed with liposomes, and power saturation ESR spectra were collected in the presence of either argon, O_2_, or NiEDDA. O_2_ and NiEDDA are spin relaxation reagents. Their concentration around the spins is correlated to their collision with the spins, and thus affects the power saturation curve of the spins (i.e., peak-to-peak amplitude vs microwave power), from which the accessibility parameters Π(O_2_) and Π(NiEDDA) are calculated (Fanucci & Cafiso, 2006; Hubbell *et al*, 1998). The CW-ESR measurement spectra were collected on an ELEXSYS ESR spectrometer at X-band (9.5 GHz) at RT. The power saturation experiments were performed in air, argon, and 20 mM Ni(II)-diammine-2,2’-(ethane-1,2-diyldiimino) diacetic acid (NiEDDA) with argon conditions. The latter two conditions were achieved by repeatedly degassing and saturating the sample with argon (Georgieva *et al.*, 2014). In each condition, the spectra were recorded as a function of microwave power, which was varied from 0.1 mW to 200 mW in 30 steps. The number of scans depended on the quality of the signal. The half-saturation parameter (P_1/2_) is obtained by fitting the equation A=I*√P* [1+(2^1/ε^-1)*P/ P_1/2_]^-ε^, where P is the microwave power applied, A is the peak-to-peak value of the central line of the spectra, and ε is the line-homogeneity parameter that we obtained from the fitting (usually ε = 1.5 formed the best fit). The accessibility parameter Π(O_2_) and Π(Ni) are calculated by the equation Π(O_2_)=[P_1/2_(O_2_)/ ∆H(O_2_) - P_1/2_(Ar)/∆H(Ar)]/[P_1/2_(ref)/∆H(ref)], and Π(Ni) = [P_1/2_(Ni)/ ∆H(Ni) - P_1/2_(Ar)/∆H(Ar)]/[P_1/2_(Ref)/∆H(Ref)], where ∆H is the line width of the central line measured at 2 mW. The insertion depth parameter Φ, which is independent of the reference, was calculated by the equation Φ=ln[Π(O_2_)/(Π(NiEDDA)] (Altenbach *et al*, 1994; Georgieva *et al.*, 2014). The P_1/2_(ref) and ∆H(ref) are typically obtained from a standard sample, 2,2-diphenyl-1-picrylhydrazyl (DPPH), to account for differences in resonator efficiencies (P_1/2_) and compensates for differences in the spin-spin relaxation time (T_2_) by factoring in the central line width (∆H) (Farahbakhsh *et al*, 1992). However, the [P_1/2_(ref)/∆H(ref)] term has been cancelled in the calculation of the insert depth parameter Φ. Therefore, neither the P_1/2_(ref) nor ∆H(ref) were used in calculations. All experiments were done at least in duplicate to ensure reproducibility. Error reported is the standard deviation.

### Small-angle X-ray scattering (SAXS)

Lyophilized phospholipids 1,2-dioleoyl-*sn*-glycero-3-phospho-L-serine (sodium salt) (DOPS) and 1,2-dioleoyl-*sn*-glycero-3-phosphoethanolamine (DOPE) were purchased from Avanti Polar Lipids and dissolved in chloroform at 20 mg/mL to produce individual lipid stock solutions. The lipid stock solutions were mixed at a molar ratio of 1/4 DOPS/DOPE, evaporated under nitrogen, and desiccated overnight under vacuum to form a dry lipid film. The lipid film was resuspended in aqueous pH 7.4 buffer (10 mM HEPES, 140 mM NaCl) to a concentration of 20 mg/mL. The resulting lipid suspension was incubated overnight at 37°C, sonicated until clear, and extruded through a 0.2 μm pore size Anotop syringe filter (Whatman) to form SUVs.

Lyophilized peptides UL31^(41-50)^, UL31^(22-42)^, and UL34^(174-194)^ were solubilized in aqueous pH 7.4 buffer (10 mM HEPES, 140 mM NaCl) and mixed with SUVs at peptide-to-lipid charge ratios (ch P/L) of 1/4, 1/2, 1/1, 3/2, and 2/1, which correspond to peptide-to-lipid molar ratios (mol P/L) of 1/140, 1/70, 1/35, 3/70, and 2/35 for UL31^(22-42)^ and UL34^(174-194)^, and 1/80, 1/40, 1/20, 3/40, and 1/10 for UL31^(41-50)^. Peptide–lipid samples were hermetically sealed into quartz capillaries (Hilgenberg GmbH, Mark-tubes, Cat. No. 4017515) and incubated at 37° C. SAXS measurements taken at the Stanford Synchrotron Radiation Lightsource (SSRL, beamline 4-2) using monochromatic X-rays with an energy of 9 keV. The scattered radiation was collected using a DECTRIS PILATUS3 × 1M detector (172 μm pixel size) and the resulting 2D SAXS powder patterns integrated using the Nika 1.82 (Ilavsky, 2012) package for Igor Pro 7.08 (WaveMetrics).

The integrated scattering intensity *I(Q)* vs. *Q* was plotted using OriginPro 2017 and the ratios of the measured peak *Q* positions were compared with those of permitted reflections for different crystal phases to identify the phase(s) present in each sample. For a cubic phase, *Q* = (2π/*a*)√(*h*^2^ + *k*^2^ + *l*^2^), and for a hexagonal phase, *Q* = (4π/(*a*√3))√(*h*^2^ + *hk* + *k*^2^), where *a* is the lattice parameter and *h*, *k*, *l* are the Miller indices of the reflection. Linear regressions of measured *Q* vs. √(*h*^2^ + *k*^2^ + *l*^2^) for cubic phases and measured *Q* vs. √(*h*^2^ + *hk* + *k*^2^) for hexagonal phases were performed. The slope *m* of each regression was then used to calculate the respective cubic (*m* = 2π/*a*) and hexagonal (*m* = 4π/(*a*√3)) lattice parameters. For a lamellar phase, the periodicity, *d*, can be calculated from the relation of *Q* = 2π*n*/*d*, where *n* is the order of the reflection.

For a cubic phase, the average Gaussian curvature per unit cell is calculated using the equation <*K*> = (2πχ)/(*A*_0_*a*^2^), where the Euler characteristic, χ, and the dimensionless surface area per unit cell, *A*_0_, are constants specific to each cubic phase (Shearman *et al*, 2006). For *Pn3m*, χ = –2 and *A*_0_ = 1.919. For *Im3m*, χ = –4 and *A*_0_ = 2.345.

### Double electron-electron resonance (DEER) spectroscopy

Approximately 50 μM peptide or peptide mixture were incubated with 10 mM SUVs (3/1/1 POPC/POPS/POPA molar ratio) in a 1/1 ratio for 10 minutes at 25 °C. Deuterated glycine was added to reach a final concentration of 20% (w/v). The samples were transferred to an ESR tube and rapidly frozen in liquid nitrogen. Standard four-pulse DEER ESR experiments were performed using a Bruker 34 GHz Q-band ELEXSYS ESR spectrometer (Bruker Instruments, Billerica, MA) at 60 K. A pulse sequence with π/2− π− π pulse widths of 16 ns, 32 ns and 32 ns, respectively and a 32 ns π pump pulse was routinely used or adjusted by the standard setup experiments. The frequency separation between detection and pump pulses was typically 70 MHz or else determined in standard setup experiments. Typical evolution times were 6 μs with signal averaging from 8-10 hours. The spectra were subject to wavelet denoising (Srivastava *et al.*, 2016) as necessary. The background signals were removed from the raw time domain signals, and the distances were reconstructed from the baseline-subtracted signals using the singular value decomposition (SVD) method (Srivastava & Freed, 2019). The P(r) distributions obtained by this method were compared to the ones using the Tikhonov regulation method and refined by the maximum entropy method as previously described (Borbat & Freed, 2007; Chiang *et al*, 2005). In our case, the difference between these two methods were not significant. The distance distribution is further fitted by a Gaussian distribution to obtain the position and width of the peak. The data were analyzed using Origin (OriginLab Inc.). Data reported is from at least two individual experiments with the error reported as the standard error of the mean.

### Chemical crosslinking

A total of 50 μM peptide(s) in PBS was incubated with or without 1 mM SUVs (3/1/1 POPC/POPS/POPA molar ratio) (<100 nm) for 10 minutes at room temperature. In crosslinking experiments all peptides were N-terminally acetylated and C-terminally amidated. In the case of two peptide mixtures, 25 μM of each peptide was used. SM(PEG)_6_ crosslinker (ThermoFisher Scientific), containing *N*-hydroxysuccinimide and maleimide groups that react with primary amines and sulfhydryls, respectively, was added at a 50-fold molar excess, and the samples were incubated for 30 minutes at room temperature. The reaction was stopped by adding Tris-HCl, pH 8.0 to a final concentration of 25 mM and glutathione to a final concentration of 50 mM. Samples were analyzed by 16% Tris-Tricine-SDS-PAGE and Coomassie staining. For each sample, band intensities of the higher molecular weight crosslinked protein were integrated and expressed as a percentage of the integrated intensity of uncrosslinked protein. Each experiment was done in duplicate, and the average value and standard error of the mean is reported.

### Circular dichroism (CD)

Far-UV CD spectra of peptides +/− SUVs were recorded using the Jasco 815 CD Spectropolarimeter at the Center for Macromolecular Interactions at Harvard Medical School. All peptides and vesicles were in 10 mM Na phosphate, pH 7.4, and 100 mM NaF buffer. Data were collected at ambient temperature with a scan speed of 50 nm/min and 5 accumulations of each sample was averaged. The raw data was background subtracted for the presence or absence vesicles and converted to mean residue ellipticity (MRE) and plotted using GraphPad Prism 9.0.

## Acknowledgements

We thank Janna Bigalke for generating plasmids pJB02, pJB41, pJB57 and pJB60 and for initiating the studies of the NEC220-SE6 mutant. We thank Elizabeth Draganova for generating the budding data for the NEC220-SE6-His_8_ mutant. We thank Alenka Lovy (Tufts University School of Medicine) for assistance with fluorescence microscopy experiments and Kelly Arnett (Harvard Medical School) for help with the circular dichroism experiments. We thank Matthew Robinson at the Center for Nanoscale Science and Technology at the National Institute of Standards and Technology, U.S. Department of Commerce, for performing the sputtered thin film depositions. We also thank Peter Cherepanov (Francis Crick Institute) for the gift of the GST-PreScission protease expression plasmid, Thomas Schwartz (Massachusetts Institute of Technology) for the gift of LoBSTr cells, and David Vanderah (Institute for Bioscience and Biotechnology Research) for the gift of the HC18 tether molecule. This work was funded by the NIH grants R01GM111795 (E.E.H.), R01AI147625 (E.E.H.), R01GM067180 (M.W.L. and G.C.L.W.), and R01GM123779 (J.H.F.), NSF grant DMR1808459 (M.W.L. and G.C.L.W.), and by a Faculty Scholar grant 55108533 from Howard Hughes Medical Institute (E.E.H.). M.K.T. was supported by the Rosenberg Fellowship (Tufts University School of Medicine). Confocal microscopy was performed at the Tufts Center for Neuroscience Research at Tufts University School of Medicine supported by NIH grant P30 NS047243 (Rob Jackson). Circular dichroism experiments were performed at the Center for Macromolecular Interactions at Harvard Medical School. ESR experiments were conducted at the National Biomedical Center for Advanced ESR Technology (ACERT), funded by NIH grant P41GM103521 (J.H.F.). SAXS experiments were conducted at the Stanford Synchrotron Radiation Lightsource (SSRL), SLAC National Accelerator Laboratory, which is supported by the U.S. Department of Energy, Office of Science, Office of Basic Energy Sciences under Contract No. DE-AC02-76SF00515. The SSRL Structural Molecular Biology Program is supported by the U.S. Department of Energy, Office of Biological and Environmental Research, and by the NIH grant P30GM133894. NR experiments were conducted at the National Institute of Standards and Technology Center for Neutron Research (NIST NCNR) on the off-specular reflectometer (MAGIK). This work used the Extreme Science and Engineering Discovery Environment (XSEDE), which is supported by National Science Foundation grant number ACI-1053575. Specifically, it used the Bridges system, which is supported by NSF award number ACI-1445606, at the Pittsburgh Supercomputing Center (PSC). Support for M.K.T to attend the Center for High Resolution Neutron Scattering Summer School on Neutron Scattering was provided by the Center for High Resolution Neutron Scattering, a partnership between the National Institute of Standards and Technology and the National Science Foundation under Agreement No. DMR-2010792. Certain commercial materials, equipment, and instruments are identified in this work to describe the experimental procedure as completely as possible. In no case does such an identification imply a recommendation or endorsement by NIST, nor does it imply that the materials, equipment, or instruments identified are necessarily the best available for the purpose.

## Author contributions

M.K.T. and E.E.H. designed and coordinated the project. M.K.T. cloned, expressed, and purified all NEC proteins and UL31 1-50 peptides as well as performed all *in vitro* budding assays, binding assays (co-sedimentation and co-flotation), sequence alignments, chemical crosslinking and circular dichroism experiments under the guidance of E.E.H. M.K.T. and D.P.H. collected NR data. D.P.H. processed the NR data. A.L.L. collected and processed ESR data. A.L.L and J.H.F. analyzed the ESR data. M.W.L. screened peptide sequences using the machine-learning classifier, performed the SAXS experiments, and analyzed the data under the guidance of G.C.L.W. All authors contributed to writing the manuscript.

## References

AlexanderNS, SteinRA, KoteicheHA, KaufmannKW, McHaourabHS, Meiler J (2013) RosettaEPR: rotamer library for spin label structure and dynamics. PloS one 8: e72851-e72851

AlonsoYAM, MiglianoSM, Teis D (2016) ESCRT-III and Vps4: a dynamic multipurpose tool for membrane budding and scission. FEBS J 283: 3288–3302

AltenbachC, GreenhalghDA, KhoranaHG, Hubbell WL (1994) A collision gradient method to determine the immersion depth of nitroxides in lipid bilayers: application to spin-labeled mutants of bacteriorhodopsin. Proceedings of the National Academy of Sciences of the United States of America 91: 1667–1671

Ames BN (1966) [10] Assay of inorganic phosphate, total phosphate and phosphatases. In: Methods in Enzymology,. 115–118. Academic Press:

AriiJ, TakeshimaK, MaruzuruY, KoyanagiN, KatoA, Kawaguchi Y (2019) Roles of the Inter-Hexamer Contact Site for Hexagonal Lattice Formation of Herpes Simplex Virus 1 Nuclear Egress Complex in Viral Primary Envelopment and Replication. Journal of Virology: JVI.00498-00419

AriiJ, WatanabeM, MaedaF, Tokai-NishizumiN, ChiharaT, MiuraM, MaruzuruY, KoyanagiN, KatoA, Kawaguchi Y (2018) ESCRT-III mediates budding across the inner nuclear membrane and regulates its integrity. Nat Commun 9: 3379

BassereauP, JinR, BaumgartT, DesernoM, DimovaR, FrolovVA, BashkirovPV, Grubmüller H, JahnR, Risselada HJ et al (2018) The 2018 biomembrane curvature and remodeling roadmap. Journal of physics D: Applied physics 51: 343001

BeurdenS, Engelsma M (2012) Herpesviruses of Fish, Amphibians and Invertebrates. In:

BigalkeJM, Heldwein EE (2015) Structural basis of membrane budding by the nuclear egress complex of herpesviruses. EMBO J 34: 2921–2936

BigalkeJM, Heldwein EE (2017) Chapter Three - Have NEC Coat, Will Travel: Structural Basis of Membrane Budding During Nuclear Egress in Herpesviruses. In: Advances in Virus Research, Margaret Kielian T.C.M., Marilyn J.R. (eds.). 107–141. Academic Press:

BigalkeJM, HeuserT, NicastroD, Heldwein EE (2014) Membrane deformation and scission by the HSV-1 nuclear egress complex. Nat Commun 5: 4131

BorbatPP, FreedJH, 2007. Measuring Distances by Pulsed Dipolar ESR Spectroscopy: Spin - Labeled Histidine Kinases, Methods in Enzymology. Elsevier,. 52–116.

BorbatPP, GeorgievaER, Freed JH (2013) Improved Sensitivity for Long-Distance Measurements in Biomolecules: Five-Pulse Double Electron-Electron Resonance. The journal of physical chemistry letters 4: 170–175

BubeckA, WagnerM, RuzsicsZ, LotzerichM, IglesiasM, SinghIR, Koszinowski UH (2004) Comprehensive mutational analysis of a herpesvirus gene in the viral genome context reveals a region essential for virus replication. J Virol 78: 8026–8035

BubeckD, FilmanDJ, Hogle JM (2005) Cryo-electron microscopy reconstruction of a poliovirus-receptor-membrane complex. Nature Structural & Molecular Biology 12: 615–618

BudilDE, LeeS, SaxenaS, Freed JH (1996) Nonlinear-Least-Squares Analysis of Slow-Motion EPR Spectra in One and Two Dimensions Using a Modified Levenberg–Marquardt Algorithm. Journal of Magnetic Resonance, Series A 120: 155–189

BuserCA, McLaughlin S (1998) Ultracentrifugation Technique for Measuring the Binding of Peptides and Proteins to Sucrose-Loaded Phospholipid Vesicles. In: Transmembrane Signaling Protocols, Bar-Sagi D. (ed.). 267–281. Humana Press: Totowa, NJ

CampeloF, FabrikantG, McMahonHT, Kozlov MM (2010) Modeling membrane shaping by proteins: focus on EHD2 and N-BAR domains. FEBS Lett 584: 1830–1839

CampeloF, McMahonHT, Kozlov MM (2008a) The Hydrophobic Insertion Mechanism of Membrane Curvature Generation by Proteins. Biophysical Journal 95: 2325–2339

CampeloF, McMahonHT, Kozlov MM (2008b) The hydrophobic insertion mechanism of membrane curvature generation by proteins. Biophys J 95: 2325–2339

ChangYE, Roizman B (1993) The product of the UL31 gene of herpes simplex virus 1 is a nuclear phosphoprotein which partitions with the nuclear matrix. J Virol 67: 6348–6356

ChangYE, Van Sant C, KrugPW, SearsAE, Roizman B (1997) The null mutant of the U(L)31 gene of herpes simplex virus 1: construction and phenotype in infected cells. Journal of virology 71: 8307–8315

ChenFY, LeeMT, Huang HW (2003) Evidence for membrane thinning effect as the mechanism for peptide-induced pore formation. Biophys J 84: 3751–3758

ChiangY-W, BorbatPP, Freed JH (2005) The determination of pair distance distributions by pulsed ESR using Tikhonov regularization. Journal of Magnetic Resonance 172: 279–295

CrumpCM, YatesC, Minson T (2007) Herpes simplex virus type 1 cytoplasmic envelopment requires functional Vps4. Journal of virology 81: 7380–7387

DasS, DixonJE, Cho W (2003) Membrane-binding and activation mechanism of PTEN. Proceedings of the National Academy of Sciences of the United States of America 100: 7491–7496

DesaiPJ, PryceEN, HensonBW, LuitweilerEM, Cothran J (2012) Reconstitution of the Kaposi’s sarcoma-associated herpesvirus nuclear egress complex and formation of nuclear membrane vesicles by coexpression of ORF67 and ORF69 gene products. J virol 86: 594–598

DraganovaEB, ThorsenMK, Heldwein EE (2020) Nuclear Egress. Curr Issues Mol Biol 41: 125–170

DuraJA, PierceDJ, MajkrzakCF, MaliszewskyjNC, McGillivrayDJ, LoscheM, O’Donovan KV, MihailescuM, Perez-SalasU, Worcester DL et al (2006) AND/R: Advanced neutron diffractometer/reflectometer for investigation of thin films and multilayers for the life sciences. Rev Sci Instrum 77: 074301

EellsR, HoogerheideDP, KienzlePA, Lösche M, MajkrzakCF, HeinrichF, 2019. 3. Structural investigations of membrane-associated proteins by neutron reflectometry, Characterization of Biological MembranesStructure and Dynamics.

EffantinG, DordorA, SandrinV, MartinelliN, SundquistWI, SchoehnG, Weissenhorn W (2013) ESCRT-III CHMP2A and CHMP3 form variable helical polymers in vitro and act synergistically during HIV-1 budding. Cellular microbiology 15: 213–226

FanucciG, Cafiso D (2006) Recent advances and applications of site-directed spin labeling. Curr Opin Struct Biol 16: 644–653

FarahbakhshZT, AltenbachC, Hubbell WL (1992) SPIN LABELED CYSTEINES AS SENSORS FOR PROTEIN-LIPID INTERACTION AND CONFORMATION IN RHODOPSIN. Photochemistry and Photobiology 56: 1019–1033

FarinaA, FeederleR, RaffaS, GonnellaR, SantarelliR, FratiL, AngeloniA, TorrisiMR, FaggioniA, Delecluse H-J (2005) BFRF1 of Epstein-Barr Virus Is Essential for Efficient Primary Viral Envelopment and Egress. Journal of Virology 79: 3703–3712

FlintJ, RacanielloVR, RallGF, Skalka AM (2015) Principles of Virology, Fourth Edition, Bundle. American Society of Microbiology

FuchsW, KluppBG, GranzowH, OsterriederN, Mettenleiter TC (2002) The Interacting UL31 and UL34 Gene Products of Pseudorabies Virus Are Involved in Egress from the Host-Cell Nucleus and Represent Components of Primary Enveloped but Not Mature Virions. Journal of Virology 76: 364–378

GeM, Freed JH (2003) Hydration, Structure, and Molecular Interactions in the Headgroup Region of Dioleoylphosphatidylcholine Bilayers: An Electron Spin Resonance Study. Biophysical Journal 85: 4023–4040

GeM, Freed JH (2009) Fusion Peptide from Influenza Hemagglutinin Increases Membrane Surface Order: An Electron-Spin Resonance Study. Biophysical Journal 96: 4925–4934

GeM, Freed JH (2011) Two Conserved Residues Are Important for Inducing Highly Ordered Membrane Domains by the Transmembrane Domain of Influenza Hemagglutinin. Biophysical Journal 100: 90–97

GeMT, Costa-FilhoA, GidwaniA, HolowkaD, BairdB, Freed J (2001) The structure of bleb membranes of RBL-2H3 cell is heterogenous: An ESR study. Biophysical Journal 80: 332a-332a

GeorgievaER, BorbatPP, NormanHD, Freed JH (2015) Mechanism of influenza A M2 transmembrane domain assembly in lipid membranes. Scientific reports 5: 11757–11757

GeorgievaER, RamlallTF, BorbatPP, FreedJH, Eliezer D (2010) The lipid-binding domain of wild type and mutant alpha-synuclein: compactness and interconversion between the broken and extended helix forms. The Journal of biological chemistry 285: 28261–28274

GeorgievaER, XiaoS, BorbatPP, FreedJH, Eliezer D (2014) Tau binds to lipid membrane surfaces via short amphipathic helices located in its microtubule-binding repeats. Biophysical journal 107: 1441–1452

GordonTB, HaywardJA, MarshGA, BakerML, Tachedjian G (2019) Host and Viral Proteins Modulating Ebola and Marburg Virus Egress. Viruses 11

GranatoM, FeederleR, FarinaA, GonnellaR, SantarelliR, HubB, FaggioniA, Delecluse H-J (2008) Deletion of Epstein-Barr Virus BFLF2 Leads to Impaired Viral DNA Packaging and Primary Egress as Well as to the Production of Defective Viral Particles. Journal of Virology 82: 4042–4051

Greenfield NJ (2006) Using circular dichroism spectra to estimate protein secondary structure. Nature Protocols 1: 2876–2890

HagenC, DentKC, Zeev-Ben-Mordehai T, GrangeM, BosseJB, WhittleC, KluppBG, SiebertCA, VasishtanD, Bauerlein FJ et al (2015) Structural Basis of Vesicle Formation at the Inner Nuclear Membrane. Cell 163: 1692–1701

HaugoAC, SzparaML, ParsonsL, EnquistLW, Roller RJ (2011) Herpes simplex virus 1 pUL34 plays a critical role in cell-to-cell spread of virus in addition to its role in virus replication. J Virol 85: 7203–7215

HubbellWL, GrossA, LangenR, Lietzow MA (1998) Recent advances in site-directed spin labeling of proteins. Curr Opin Struct Biol 8: 649–656

Hurley JH (2015) ESCRTs are everywhere. EMBO J 34: 2398–2407

HurleyJH, Cada AK (2018) Inside job: how the ESCRTs release HIV-1 from infected cells. Biochem Soc Trans 46: 1029–1036

Ilavsky J (2012) Nika: software for two-dimensional data reduction. Journal of Applied Crystallography 45: 324–328

ItohT, ErdmannKS, RouxA, HabermannB, WernerH, De Camilli P (2005) Dynamin and the Actin Cytoskeleton Cooperatively Regulate Plasma Membrane Invagination by BAR and F-BAR Proteins. Developmental Cell 9: 791–804

JohnsonDC, Baines JD (2011) Herpesviruses remodel host membranes for virus egress. Nature reviews Microbiology 9: 382–394

KaplanA, LeeMW, WolfAJ, LimonJJ, BeckerCA, DingM, MuraliR, LeeEY, LiuGY, Wong GCL et al (2017) Direct Antimicrobial Activity of IFN-β. The Journal of Immunology 198: 4036–4045

KatoA, YamamotoM, OhnoT, KodairaH, NishiyamaY, Kawaguchi Y (2005) Identification of proteins phosphorylated directly by the Us3 protein kinase encoded by herpes simplex virus 1. Journal of virology 79: 9325–9331

KeennaTW, BerezneyR, FunkLK, Crane FL (1970) Lipid composition of nuclear membranes isolated from bovine liver. Biochimica et Biophysica Acta (BBA) - Biomembranes 203: 547–554

KellySM, JessTJ, Price NC (2005) How to study proteins by circular dichroism. Biochimica et Biophysica Acta (BBA) - Proteins and Proteomics 1751: 119–139

KienzlePA, KryckaJ, PatelN, MettingC, SahinI, FuZ, ChenW, MontA, TigheD, 2016. Refl1D (Version 0.7.7) [Computer Software]. College Park, MD: University of Maryland.

Kim J, BlackshearPJ, JohnsonJD, McLaughlin S (1994a) Phosphorylation reverses the membrane association of peptides that correspond to the basic domains of MARCKS and neuromodulin. Biophysical journal 67: 227–237

KimJ, ShishidoT, XlJ, AderemA, McLaughlin S (1994b) Phosphorylation, high ionic strength, and calmodulin reverse the binding of MARCKS to phospholipid vesicles. The Journal of biological chemistry 269: 28214–28219

KimYW, Sung W (2001) Membrane curvature induced by polymer adsorption. Physical Review E 63: 041910

KirbyBJ, KienzlePA, MaranvilleBB, BerkNF, KryckaJ, HeinrichF, Majkrzak CF (2012) Phase-sensitive specular neutron reflectometry for imaging the nanometer scale composition depth profile of thin-film materials. Curr Opin Colloid Int Sci 17: 44–53

KluppBG, GranzowH, FuchsW, KeilGM, FinkeS, Mettenleiter TC (2007) Vesicle formation from the nuclear membrane is induced by coexpression of two conserved herpesvirus proteins. Proc Natl Acad Sci U S A 104: 7241–7246

KluppBG, GranzowH, Mettenleiter TC (2000) Primary Envelopment of Pseudorabies Virus at the Nuclear Membrane Requires the UL34 Gene Product. Journal of Virology 74: 10063–10073

KluppBG, HellbergT, Rönfeldt S, FranzkeK, FuchsW, Mettenleiter TC (2018) Function of the non-conserved N-terminal domain of Pseudorabies Virus pUL31 in nuclear egress. Journal of Virology

KozlovskyY, Kozlov MM (2003) Membrane Fission: Model for Intermediate Structures. Biophysical Journal 85: 85–96

KuoW, HerrickDZ, Cafiso DS (2011) Phosphatidylinositol 4,5-bisphosphate alters synaptotagmin 1 membrane docking and drives opposing bilayers closer together. Biochemistry 50: 2633–2641

Lai Alex L, Freed Jack H (2014) HIV gp41 Fusion Peptide Increases Membrane Ordering in a Cholesterol-Dependent Fashion. Biophysical Journal 106: 172–181

LaiAL, Freed JH (2015) The Interaction between Influenza HA Fusion Peptide and Transmembrane Domain Affects Membrane Structure. Biophysical journal 109: 2523–2536

LaiAL, MilletJK, DanielS, FreedJH, Whittaker GR (2017) The SARS-CoV Fusion Peptide Forms an Extended Bipartite Fusion Platform that Perturbs Membrane Order in a Calcium-Dependent Manner. J Mol Biol 429: 3875–3892

LeeEY, FulanBM, WongGC, Ferguson AL (2016) Mapping membrane activity in undiscovered peptide sequence space using machine learning. Proc Natl Acad Sci U S A 113: 13588–13593

LeeMW, LeeEY, LaiGH, KennedyNW, PoseyAE, XianW, FergusonAL, HillRB, Wong GCL (2017) Molecular Motor Dnm1 Synergistically Induces Membrane Curvature To Facilitate Mitochondrial Fission. ACS Cent Sci 3: 1156–1167

LiangZC, Freed JH (1999) An assessment of the applicability of multifrequency ESR to study the complex dynamics of biomolecules. Journal of Physical Chemistry B 103: 6384–6396

LicataJM, Simpson-HolleyM, WrightNT, HanZ, ParagasJ, Harty RN (2003) Overlapping motifs (PTAP and PPEY) within the Ebola virus VP40 protein function independently as late budding domains: involvement of host proteins TSG101 and VPS-4. J Virol 77: 1812–1819

LorenzM, VollmerB, UnsayJD, KluppBG, Garcia-Saez AJ, MettenleiterTC, Antonin W (2015a) A Single Herpesvirus Protein Can Mediate Vesicle Formation in the Nuclear Envelope. Journal of Biological Chemistry 290: 6962–6974

LorenzM, VollmerB, UnsayJD, KluppBG, Garcia-Saez AJ, MettenleiterTC, Antonin W (2015b) A single herpesvirus protein can mediate vesicle formation in the nuclear envelope. J Biol Chem 290: 6962–6974

Lötzerich M, RuzsicsZ, Koszinowski UH (2006) Functional domains of murine cytomegalovirus nuclear egress protein M53/p38. Journal of virology 80: 73–84

LyeMF, SharmaM, El Omari K, FilmanDJ, SchuermannJP, HogleJM, Coen DM (2015) Unexpected features and mechanism of heterodimer formation of a herpesvirus nuclear egress complex. EMBO J 34: 2937–2952

McCulloughJ, FrostA, Sundquist WI (2018) Structures, Functions, and Dynamics of ESCRT-III/Vps4 Membrane Remodeling and Fission Complexes. Annu Rev Cell Dev Biol 34: 85–109

McMahonHT, Boucrot E (2015) Membrane curvature at a glance. J Cell Sci 128: 1065–1070

Mettenleiter TC (2016) Breaching the Barrier—The Nuclear Envelope in Virus Infection. J Mol Biol 428: 1949–1961

MettenleiterTC, KluppBG, Granzow H (2009) Herpesvirus assembly: An update. Virus Research 143: 222–234

MihailescuM, KrepkiyD, MilescuM, GawrischK, SwartzKJ, White S (2014) Structural interactions of a voltage sensor toxin with lipid membranes. Proc Natl Acad Sci U S A 111: E5463-5470

MihailescuM, SorciM, SeckuteJ, SilinVI, HammerJ, PerrinBSJr., HernandezJI, SmajicN, ShresthaA, Bogardus KA et al (2019) Structure and Function in Antimicrobial Piscidins: Histidine Position, Directionality of Membrane Insertion, and pH-Dependent Permeabilization. J Am Chem Soc 141: 9837–9853

MishraA, LaiGH, SchmidtNW, SunVZ, RodriguezAR, TongR, TangL, ChengJ, DemingTJ, Kamei DT et al (2011) Translocation of HIV TAT peptide and analogues induced by multiplexed membrane and cytoskeletal interactions. Proceedings of the National Academy of Sciences 108: 16883–16888

Modis Y (2014) Relating structure to evolution in class II viral membrane fusion proteins. Curr Opin Virol 5: 34–41

MouF, WillsE, Baines JD (2009) Phosphorylation of the U(L)31 Protein of Herpes Simplex Virus 1 by the U(S)3-Encoded Kinase Regulates Localization of the Nuclear Envelopment Complex and Egress of Nucleocapsids. Journal of Virology 83: 5181–5191

Mulgrew-NesbittA, DiraviyamK, WangJ, SinghS, MurrayP, LiZ, RogersL, MirkovicN, Murray D (2006) The role of electrostatics in protein-membrane interactions. Biochim Biophys Acta 1761: 812–826

NathanL, LaiAL, MilletJK, StrausMR, FreedJH, WhittakerGR, Daniel S (2020) Calcium Ions Directly Interact with the Ebola Virus Fusion Peptide To Promote Structure-Function Changes That Enhance Infection. ACS Infect Dis 6: 250–260

NeitchevaT, Peeva D (1995) Phospholipid composition, phospholipase A2 and sphingomyelinase activities in rat liver nuclear membrane and matrix. The International Journal of Biochemistry & Cell Biology 27: 995–1001

NeubauerA, RudolphJ, Brandmüller C, Just FT, Osterrieder N (2002) The Equine Herpesvirus 1 UL34 Gene Product Is Involved in an Early Step in Virus Egress and Can Be Efficiently Replaced by a UL34-GFP Fusion Protein. Virology 300: 189–204

NewcombWW, FontanaJ, WinklerDC, ChengN, HeymannJB, Steven AC (2017) The Primary Enveloped Virion of Herpes Simplex Virus 1: Its Role in Nuclear Egress. MBio 8

NguyenHC, TalledgeN, McCulloughJ, SharmaA, MossFR3rd, IwasaJH, VershininMD, SundquistWI, Frost A (2020) Membrane constriction and thinning by sequential ESCRT-III polymerization. Nat Struct Mol Biol 27: 392–399

NystromNA, LevineMJ, RoskiesRZ, ScottJR, 2015. Bridges, Proceedings of the 2015 XSEDE Conference on Scientific Advancements Enabled by Enhanced Cyberinfrastructure - XSEDE ‘15. ACM, St. Louis, Missouri,. 1–8.

PeterBJ, KentHM, MillsIG, VallisY, ButlerPJG, EvansPR, McMahon HT (2004) BAR Domains as Sensors of Membrane Curvature: The Amphiphysin BAR Structure. Science 303: 495–499

PinelloJF, LaiAL, MilletJK, Cassidy-HanleyD, FreedJH, Clark TG (2017) Structure-Function Studies Link Class II Viral Fusogens with the Ancestral Gamete Fusion Protein HAP2. Curr Biol 27: 651–660

PowellKA, ValovaVA, MalladiCS, JensenON, LarsenMR, Robinson PJ (2000) Phosphorylation of Dynamin I on Ser-795 by Protein Kinase C Blocks Its Association with Phospholipids. Journal of Biological Chemistry 275: 11610–11617

QuanA, XueJ, WielensJ, SmillieKJ, AnggonoV, ParkerMW, CousinMA, GrahamME, Robinson PJ (2012) Phosphorylation of syndapin I F-BAR domain at two helix-capping motifs regulates membrane tubulation. Proceedings of the National Academy of Sciences of the United States of America 109: 3760–3765

RakovskaB, RagaliauskasT, MickeviciusM, JankunecM, NiauraG, VanderahDJ, Valincius G (2015) Structure and function of the membrane anchoring self-assembled monolayers. Langmuir 31: 846–857

Roberts-Galbraith RH, OhiMD, BallifBA, ChenJ-S, McLeodI, McDonaldWH, GygiSP, YatesJR3rd, Gould KL (2010) Dephosphorylation of F-BAR protein Cdc15 modulates its conformation and stimulates its scaffolding activity at the cell division site. Molecular cell 39: 86–99

Roizman PEPaB (2013) Herpesviridae. In: Fields Virology, David M. Knipe P.H. (ed.)Wolters Kluwer Lippincott Williams & Wilkins:

RollerRJ, Baines JD (2017) Herpesvirus Nuclear Egress. In: Cell Biology of Herpes Viruses, Osterrieder K. (ed.). 143–169. Springer International Publishing: Cham

RollerRJ, BjerkeSL, HaugoAC, Hanson S (2010) Analysis of a charge cluster mutation of herpes simplex virus type 1 UL34 and its extragenic suppressor suggests a novel interaction between pUL34 and pUL31 that is necessary for membrane curvature around capsids. J Virol 84: 3921–3934

RollerRJ, ZhouY, SchnetzerR, FergusonJ, DeSalvo D (2000) Herpes Simplex Virus Type 1 UL34 Gene Product Is Required for Viral Envelopment. Journal of Virology 74: 117–129

RossmanJS, JingX, LeserGP, Lamb RA (2010) Influenza virus M2 protein mediates ESCRT-independent membrane scission. Cell 142: 902–913

RossmanJS, Lamb RA (2011) Influenza virus assembly and budding. Virology 411: 229–236

RossmanJS, Lamb RA (2013) Viral membrane scission. Annu Rev Cell Dev Biol 29: 551–569

SchmidtNW, MishraA, LaiGH, DavisM, SandersLK, TranD, GarciaA, TaiKP, McCrayPB, Ouellette AJ et al (2011) Criterion for Amino Acid Composition of Defensins and Antimicrobial Peptides Based on Geometry of Membrane Destabilization. Journal of the American Chemical Society 133: 6720–6727

SchmidtNW, MishraA, WangJ, DeGradoWF, Wong GC (2013a) Influenza virus A M2 protein generates negative Gaussian membrane curvature necessary for budding and scission. J Am Chem Soc 135: 13710–13719

SchmidtNW, MishraA, WangJ, DeGradoWF, Wong GCL (2013b) Influenza Virus A M2 Protein Generates Negative Gaussian Membrane Curvature Necessary for Budding and Scission. Journal of the American Chemical Society 135: 13710–13719

SchurFK, ObrM, HagenWJ, WanW, JakobiAJ, KirkpatrickJM, SachseC, KrausslichHG, Briggs JA (2016) An atomic model of HIV-1 capsid-SP1 reveals structures regulating assembly and maturation. Science 353: 506–508

SchusterF, KluppBG, GranzowH, Mettenleiter TC (2012) Structural determinants for nuclear envelope localization and function of pseudorabies virus pUL34. J Virol 86: 2079–2088

ShearmanGC, CesO, TemplerRH, Seddon JM (2006) Inverse lyotropic phases of lipids and membrane curvature. Journal of Physics: Condensed Matter 18: S1105-S1124

ShekharP, NandaH, LoscheM, Heinrich F (2011) Continuous distribution model for the investigation of complex molecular architectures near interfaces with scattering techniques. J Appl Phys 110: 102216–10221612

SigalCT, ZhouW, BuserCA, McLaughlinS, Resh MD (1994) Amino-terminal basic residues of Src mediate membrane binding through electrostatic interaction with acidic phospholipids. Proceedings of the National Academy of Sciences of the United States of America 91: 12253–12257

SilvermanJL, GreeneNG, KingDS, Heldwein EE (2012) Membrane requirement for folding of the herpes simplex virus 1 gB cytodomain suggests a unique mechanism of fusion regulation. Journal of virology 86: 8171–8184

SilvestriLS, RuthelG, KallstromG, WarfieldKL, SwensonDL, NelleT, IversenPL, BavariS, Aman MJ (2007) Involvement of vacuolar protein sorting pathway in Ebola virus release independent of TSG101 interaction. The Journal of infectious diseases 196 Suppl 2: S264-270

SimunovicM, EvergrenE, Callan-JonesA, Bassereau P (2019) Curving Cells Inside and Out: Roles of BAR Domain Proteins in Membrane Shaping and Its Cellular Implications. Annu Rev Cell Dev Biol 35: 111–129

SmithAK, Freed JH (2009) Determination of tie-line fields for coexisting lipid phases: an ESR study. J Phys Chem B 113: 3957–3971

SneadD, LaiAL, WraggRT, ParisottoDA, RamlallTF, DittmanJS, FreedJH, Eliezer D (2017) Unique Structural Features of Membrane-Bound C-Terminal Domain Motifs Modulate Complexin Inhibitory Function. Front Mol Neurosci 10: 154–154

SoniSP, Stahelin RV (2014) The Ebola virus matrix protein VP40 selectively induces vesiculation from phosphatidylserine-enriched membranes. J Biol Chem 289: 33590–33597

SrivastavaM, AndersonCL, Freed JH (2016) A New Wavelet Denoising Method for Selecting Decomposition Levels and Noise Thresholds. IEEE Access 4: 3862–3877

SrivastavaM, Freed JH (2017) Singular Value Decomposition Method to Determine Distance Distributions in Pulsed Dipolar Electron Spin Resonance. The journal of physical chemistry letters 8: 5648–5655

SrivastavaM, Freed JH (2019) Singular Value Decomposition Method To Determine Distance Distributions in Pulsed Dipolar Electron Spin Resonance: II. Estimating Uncertainty. The Journal of Physical Chemistry A 123: 359–370

SrivastavaM, GeorgievaER, Freed JH (2017) A New Wavelet Denoising Method for Experimental Time-Domain Signals: Pulsed Dipolar Electron Spin Resonance. J Phys Chem A 121: 2452–2465

StachowiakJC, BrodskyFM, Miller EA (2013) A cost–benefit analysis of the physical mechanisms of membrane curvature. Nature Cell Biology 15: 1019–1027

StrausMR, TangT, LaiAL, FlegelA, BidonM, FreedJH, DanielS, WhittakerGR, 2019. Ca2+ ions promote fusion of Middle East Respiratory Syndrome coronavirus with host cells and increase infectivity. Cold Spring Harbor Laboratory.

Studier FW (2005) Protein production by auto-induction in high-density shaking cultures. Protein Expression and Purification 41: 207–234

TownsJ, CockerillT, DahanM, FosterI, GaitherK, GrimshawA, HazlewoodV, LathropS, LifkaD, Peterson GD et al (2014) XSEDE: Accelerating Scientific Discovery. Computing in Science & Engineering 16: 62–74

VanegasJM, HeinrichF, RogersDM, CarsonBD, La Bauve S, VernonBC, AkgunB, SatijaS, ZhengA, Kielian M et al (2018) Insertion of Dengue E into lipid bilayers studied by neutron reflectivity and molecular dynamics simulations. Biochimica et Biophysica Acta (BBA) - Biomembranes 1860: 1216–1230

VottelerJ, Sundquist WI (2013) Virus Budding and the ESCRT Pathway. Cell Host Microbe 14: 232–241

VrugtJA, ter Braak CJF, DiksCGH, RobinsonBA, HymanJM, Higdon D (2009) Accelerating Markov Chain Monte Carlo Simulation by Differential Evolution with Self-Adaptive Randomized Subspace Sampling. Int J Nonlin Sci Num 10: 273–290

WalzerSA, Egerer-SieberC, StichtH, SevvanaM, HohlK, MilbradtJ, MullerYA, Marschall M (2015) Crystal Structure of the Human Cytomegalovirus pUL50-pUL53 Core Nuclear Egress Complex Provides Insight into a Unique Assembly Scaffold for Virus-Host Protein Interactions. J Biol Chem 290: 27452–27458

WhittakerGR, Helenius A (1998) Nuclear import and export of viruses and virus genomes. Virology 246: 1–23

YaoH, LeeMW, WaringAJ, WongGCL, Hong M (2015) Viral fusion protein transmembrane domain adopts β-strand structure to facilitate membrane topological changes for virus–cell fusion. Proceedings of the National Academy of Sciences 112: 10926–10931

Zeev-Ben-Mordehai T, WeberrussM, LorenzM, CheleskiJ, HellbergT, WhittleC, El Omari K, VasishtanD, DentKC, Harlos K et al (2015) Crystal Structure of the Herpesvirus Nuclear Egress Complex Provides Insights into Inner Nuclear Membrane Remodeling. Cell Rep 13: 2645–2652

ZemelA, Ben-ShaulA, May S (2008) Modulation of the Spontaneous Curvature and Bending Rigidity of Lipid Membranes by Interfacially Adsorbed Amphipathic Peptides. The Journal of Physical Chemistry B 112: 6988–6996

ZhaoH, HakalaM, Lappalainen P (2010) ADF/Cofilin Binds Phosphoinositides in a Multivalent Manner to Act as a PIP2-Density Sensor. Biophysical Journal 98: 2327–2336

ZhouW, ParentLJ, WillsJW, Resh MD (1994) Identification of a membrane-binding domain within the amino-terminal region of human immunodeficiency virus type 1 Gag protein which interacts with acidic phospholipids. J Virol 68: 2556–2569

ZimmerbergJ, Kozlov MM (2006) How proteins produce cellular membrane curvature. Nat Rev Mol Cell Biol 7: 9–19

